# EEG reveals how space acts as a late heuristic of time

**DOI:** 10.1101/2025.05.28.656560

**Authors:** Fabrizio Doricchi, Gabriele Scozia, Mario Pinto, Silvana Lozito, Sara Lo Presti, Mariella Pazzaglia, Massimo Silvetti, Stefano Lasaponara

**Affiliations:** Dipartimento di Psicologia - “Sapienza” Università di Roma, Via dei Marsi 78, 00185 Roma Italy; IRCCS Fondazione Santa Lucia, via Ardeatina 354, Roma Italy; Computational and Translational Neuroscience Lab (CTNLab), Institute of Cognitive Sciences and Technologies, National Research Council (CNR), Rome, Italy

**Author notes:** Corresponding author: Prof. Fabrizio Doricchi Address: Dipartimento di Psicologia - Facolta’ di Medicina e Psicologia, Sapienza Università di Roma - Via dei Marsi 78, 00185 Rome, Italy.

## Abstract

To compensate for its sensory intangibility, humans often rely on spatial metaphors, gestures, and visual tools to represent the passage of time. These spatial tools, i.e. heuristics, range from everyday practices—such as directional hand gestures to indicate past or future events—to more abstract scientific conceptualizations, such as the “curving of space-time” in the theory of relativity. Despite this widespread spatialization of time, it remains unclear to what extent space is an inherent component of the brain’s representation of time and its role in monitoring temporal durations. Here, we combine EEG-behavioral methods and neural network models of optimal decision-making to show that space is a late compensatory mechanism of time representation recruited when faster and non-spatial timekeeping mechanisms are suboptimally engaged. EEG analyses reveal a cascade-like process: spatial engagement in timekeeping follows the insufficient non-spatial encoding of time intervals, leading to delayed decisions on their length and slower response selection. Computational modelling further indicates that trial-by-trial fluctuations in the spatialization of time are explained by stochastic variations in the activity of the dopaminergic/noradrenergic (DA/NE) system and its interaction with the anterior cingulate cortex. These findings provide the first clear evidence of when, why, and how the brain recruits spatial mechanisms in the service of temporal processing and demonstrate that non-spatial and spatial timekeeping systems can be dissociated at both behavioural and electrophysiological levels.

## Introduction

Humans use spatial metaphors, gestures and visual tools to represent and communicate the flow of time. These spatial representations are rooted in biological constraints and culturally based sensorimotor habits. As forward-walking organisms, humans display an almost universal preference to represent the past as being "*left behind* " and the future as "*waiting ahead*" of their body position (Bonato et al., 2012; Casasanto & Boroditsky, 2008; Núñez & Sweetser, 2006; Teghil et al., 2021). In addition, experiences of time spatialisation linked to reading and inspection habits also mould a corresponding spatial representation of elapsing time (Pitt & Casasanto, 2020). For example, people from left-to-right reading cultures mentally place short time durations and past events on the left side of space while long durations and future events are on the right side. Vice-versa, people from right-to-left reading cultures adopt a right-to-left directional representation of time (Boroditsky et al., 2011; Callizo-Romero et al., 2020; Fuhrman et al., 2011; Ouellet et al., 2010; Pitt & Casasanto, 2020; Vallesi et al., 2014).

The major experimental evidence for the spatial representation of time durations in humans comes from the STEARC effect (Space Time Association of Response Codes; (Conson et al., 2008; Ishihara et al., 2008; Vallesi et al., 2008). The STEARC shows that in left-to-right reading participants, deciding that a time interval is “short” is faster when motor responses are released in the left side of space and deciding that a time interval is “long” is faster with responses on the right side of space, i.e., Compatible condition. In contrast, decisions are slower when “short” intervals are associated with left-side motor responses and “long” intervals with right-side ones, i.e., Incompatible condition. This effect of spatial compatibility between the duration of time intervals and the position of motor responses has suggested that time mentally unfolds in the same direction as culturally acquired inspection and reading habits. The STEARC has been advocated for pointing at an intrinsic spatial component in brain mechanisms that helps with timekeeping. Nonetheless, at variance with this widespread view, we have recently demonstrated that the STEARC is eventually found when an observer’s decision on time duration is slow, though not when it is fast (Scozia et al., 2023). In addition, the emergence of the STEARC at slower reaction times (RTs) is matched with reduced accuracy in decisions (Scozia et al., 2023). These findings indicate that faster timing judgements are not taken at the expense of timing precision. These observations show that the spatial coding of time gets progressively superimposed on faster and equally or even more accurate non-spatial mechanisms of time coding. This suggests, in turn, that studying the STEARC as a function of the speed of interval timing makes it possible to empirically separate the behavioural manifestations and neural correlates of the non-spatial and the spatial mechanisms of time coding. Here, we report the results of a new EEG study that, for the first time, investigates these brain correlates.

Previous EEG studies (see Vallesi et al., 2011) have identified the neural correlate of the STEARC in the inter-hemispheric competition underlying the selection of compatible versus incompatible manual responses to short and long durations. Specifically, Vallesi and colleagues demonstrated that, compared to incompatible mappings (left–long, right–short), compatible mappings (left–short, right–long) were associated with a larger Lateralized Readiness Potential (LRP), that is with a larger activation of the motor cortex controlling the responding hand over the motor cortex controlling the non-responding hand. In the case of the STEARC, the study of the LRP revealed an initial activation of the right motor cortex around the expected onset of short durations, indicating early activation of the left-hand response. This motor activation gradually shifted toward the left hemisphere as time elapsed, signalling the delayed activation of the right-hand response for long durations. Notably, when left-hand “incompatible” responses were associated with “long” durations, this pattern of LRP activity was not present, that is, there was no pre-activation of the right-hand response around the expected onset of short durations. Building on this evidence, the present study investigated whether variations in the strength and presence of the STEARC effect as a function of response speed are reflected in corresponding variations in LRP amplitude. For instance, one may predict that a fully developed STEARC effect during slower responses would be accompanied by a more pronounced LRP. More important, we aimed to examine whether the emergence of the STEARC effect at slower response times is preceded by modulations in EEG components associated with temporal encoding—before motor selection processes are engaged. These early EEG components, occurring around or immediately after the offset of the time interval, are unlikely to be influenced by stimulus-response compatibility effects, as decisions regarding motor output are either not yet made or are just beginning. Nonetheless, changes in the amplitude and latency of these time-encoding components as a function of subsequent RTs may reveal early neural precursors of the STEARC effect observed in trials with slow responses. To this end, we examined the EEG correlates of all sequential sensory and cognitive processes involved in the encoding of time intervals and in the subsequent selection of motor responses used to classify them as short or long. This approach aimed to reveal not only the EEG modulations that accompany slow RT trials—when the STEARC effect tends to emerge—but also the processing stage at which EEG signatures of spatial compatibility between time duration and the position of the motor response first arise. Finally, we propose a neuro-computational account of the mechanisms governing the transition from non-spatial to spatial representations of time, as supported by the present findings.

## Materials and Methods

### Participants

The number of participants was determined through an a priori power analysis (G∗Power, Faul et al., 2007) based on the effect size f(U) = .568 gathered from a previous study by (Vallesi et al., 2008). This analysis showed that 28 participants would be needed to obtain a power of .80 when employing a two-sided .05 alpha criterion of statistical significance for repeated measures within factors ANOVAs. Therefore, we tested a sample of 30 healthy adult volunteers (22 F and 8 M; mean age: 23.3 y, d.s.: 2.6). All participants were right-handed (Edinburgh Handedness Inventory > 80%), had a normal or corrected-to-normal vision and left-to-right reading style. No participant reported past or present neurological or psychiatric conditions. After data acquisition, one participant had to be excluded from further analysis due to a large number of EEG artefacts. The final sample included 29 participants (21 F and 8 M; mean age: 23.3 y, d.s.: 2.7).

### Experimental procedure and tasks

#### Apparatus and stimuli

The experiment was administered through the E-Prime software (v.2.2). Each trial began with the 500 ms presentation of a white central fixation cross (2.3° × 2.3°). A white dot target (diameter = 2.3°) replaced the fixation cross at the centre of the screen as a timing stimulus. This stimulus was presented for a short 1000 ms or a long 3000 ms interval. At the end of the target period, 1500 ms were allowed for speeded response. The inter-trial interval (ITI) started after the participant’s response and lasted 500 to 700 ms (see **Fig. 1**). Participants were asked to judge, as quickly as possible, the short or long duration of target time intervals by pressing on a computer keyboard, the “X” left-side button with the left index finger or the “M” right-side button with the right index finger (see **Fig. 1**). In a series of four blocks of trials, short intervals were associated with the left-side button and long intervals with the right-side button (Compatible condition: Comp). In another series of four blocks of trials, the association between the response side and time duration was reversed (Incompatible condition: Incomp). The order of administration of the series Comp and Incomp blocks was counterbalanced among participants. Each block consisted of 128 trials, 64 for each time interval duration. Both in the Compatible and Incompatible condition 64 trials were administered in each of the RTs bins considered in the behavioural and EEG analyses (see below). A training block, including 20 trials, was administered before each experimental block.

**Fig. 1:**
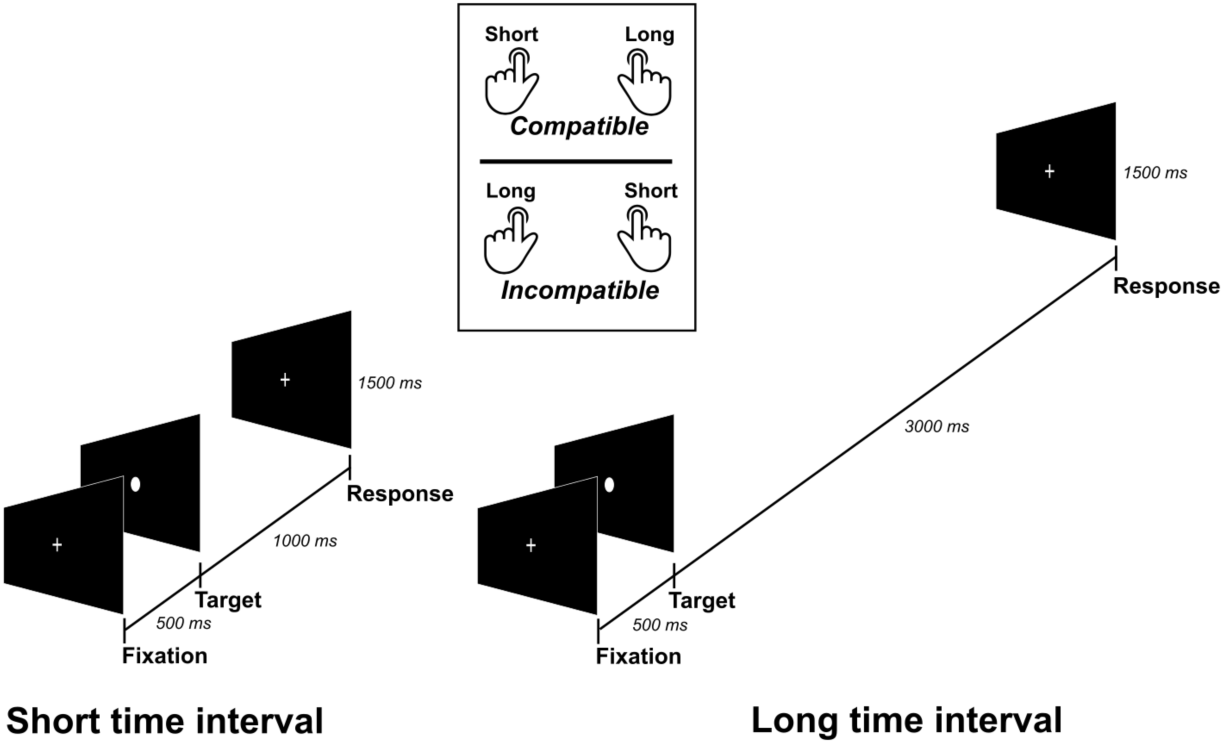
Trial events and response conditions in the STEARC task.

#### Procedure

Participants were tested in an electrically shielded, quiet and isolated room. The head position was restrained with a chin rest at a fixed viewing distance of 57.2 cm from the screen. All participants were naive to the specific aim of the study and provided written informed consent to participate in the experiment. Experimental procedures were designed following the principles of the Declaration of Helsinki and were approved before data collection by the Local Ethical Committee of “Sapienza” University of Rome (Prot. n. 0002619).

### Electrophysiological recording

The EEG was recorded from 64 electrodes placed according to the 10–10 system montage (BrainVision system). All scalp channels were referenced online to Cz. Horizontal eye movements were also monitored with a bipolar recording from electrodes at the left and right outer canthi. Blinks and vertical eye movements were recorded with an electrode below the left eye, referenced offline to site Fp1.

The EEG from each electrode site was digitised at 256 Hz with an amplifier bandpass of 0.1–1000 Hz, including a 50 Hz notch filter, and was stored for offline averaging. The EEG was initially re-referenced against the average of all channels and then filtered offline with a bandpass of 0.1–30 Hz. Successively, for the analyses locked to target-onset, continuous EEG was segmented in epochs of [-200, +1000 ms] for short intervals and [-200, +3000 ms] for long intervals. In the analyses locked to target-offset, epochs lasted [-200, +1500 ms] for short and long intervals. The 200 ms period preceding target onset or offset was used for baseline correction in all cases. Finally, EEG was segmented in epochs lasting [-1200, +300 ms] for the analysis locked to manual response. Also in this case, the first 200 ms of each epoch were used for baseline correction.

Prior to computerised artefact rejection, oculo-motor correction was performed using Independent Component Analysis (ICA) as implemented in BrainVision Analyzer 2 software. Artefact rejection was then performed before signal averaging to discard epochs containing maximum amplitudes exceeding ±100 uV and voltage steps of more than 60 μV/ms. On average, 3.4% of segments were rejected for violating these criteria.

For target onset/offset analyses, segments were averaged according to the combination of Space-Time Compatibility (Comp vs Incomp), Time interval (Short vs Long) and RTs–Bin (B1, B2, B3, B4). For response-locked analyses, segments were averaged according to the combination of manual response (Left, Right), Space–Time Compatibility (Comp vs Incomp), and RTs–Bin (B1, B2, B3, B4).

### Statistical analyses

#### Manual Reaction Times (RTs)

RTs below or above two sd from individual average RTs were excluded from analyses. Following this criterion, less than 3% of trials. The STEARC was measured by comparing the Reaction Times (RTs) advantage in the Comp versus the Incomp Condition. To investigate its temporal development, we measured the STEARC as a function of RTs speed along four quartiles/Bins ranging from the fastest RTs in Bin1 (B1) to the slowest RTs in Bin4 (B4; (Rubichi et al., 1997). To this aim, we used the Vincentization procedure (Ratcliff, 1979; see Pinto, Pellegrino, Marson, et al., 2021; Rubichi et al., 1997; Scozia et al., 2023). For each participant, we calculated the RT distributions of correct responses, i.e., from fastest to slowest, in both Comp and Incomp experimental conditions. We then divided each distribution into four proportional quartile-Bins so that each Bin contained the same proportion of trials, i.e., one-fourth of trials. The difference between mean RTs from corresponding Bins in Comp and Incomp conditions provides a Bin-by-Bin measure of the time course of the STEARC.

Finally, individual RTs were then entered in a 2 × 2 × 4 repeated-measures ANOVA to test the effect of Space-Time Compatibility (Comp vs Incomp), Time interval (Short vs Long) and RTs–Bin (B1, B2, B3, B4) as within factors. The distribution of errors was analysed through RTs–Bin (B1, B2, B3, B4) x Space–Time Compatibility (Comp vs Incomp) x Time interval (Short vs Long repeated measures ANOVA. Significant main effects and interactions were explored in all cases through Bonferroni post-hoc comparisons.

### Electrophysiological data

#### Analyses locked to target-onset: CNV component

In line with previous studies (Brunia & van Boxtel, 2000; Kononowicz & Penney, 2016; Kononowicz & Van Rijn, 2014; Ng et al., 2011; Tarantino et al., 2010), the time-related Contingent Negative Variation (CNV) was measured as mean activity over a pool of fronto-central derivations including FC1, FC2, FCz, C1, Cz and C2. Early and late phases of the CNV were of interest (see Results). An early phase late phase within the 300–550 msec time window and late phase within the 550-900 msec time windows of the CNV were found both for short and long intervals. A later phase related to long intervals was found within a 1700– 2900 msec time window. Individual mean amplitudes were entered in a Space-Time Compatibility (Comp vs Incomp) x Time interval (Short vs Long) x RTs–Bin (B1, B2, B3, B4) repeated measures ANOVAs. Significant main effects and interactions were explored in all cases through Bonferroni post-hoc comparisons.

#### Analyses locked to target-onset: Lateralized Readiness Potential (C3 - C4)

In line with a previous study by Vallesi et al., (2011), for each of our experimental conditions, i.e., Space-Time Compatibility (Comp vs Incomp), Time Interval (Short vs Long) and RTs–Bin (B1, B2, B3, B4), we calculated the differential activation of the scalp regions over the left versus right hand-motor cortex, by subtracting C4 from C3 amplitudes, in a time window of -200 to 1000 ms for Short intervals, and -200 to 3000 ms for Long ones. Since preparing a hand movement induces motor negativity over the hand-motor cortex in the hemisphere contralateral to the hand (Brunia & van Boxtel, 2000; Shibasaki et al., 1981; Toma et al., 2002), a positive C3-C4 differential waveform indicates a left-more-than-right hand motor activation, while a negative differential waveform indicates an opposite right-more-than-left hand motor activation. We evaluated the statistical significance of these differential waveforms using a series of parametric continuous two-tailed one-sample T-tests (against zero) corrected for multiple comparisons (FDR) in the time domain, as implemented in Brainstorm (Tadel et al., 2011), documented and freely available for download online under the GNU general public license at http://neuroimage.usc.edu/brainstorm).

*Analyses locked to target-offset: N1/P2, P300, and LPCt components*.

The N1/P2 complex was measured over a pool of fronto-central derivations, including FC1, FCz, and FC2 (Kononowicz & Van Rijn, 2014; Tarantino et al., 2010), during time windows of 90-180 ms and 180-290 ms, the N1 and the P2 component, respectively. For each component, individual mean amplitudes were then entered in a Space-Time Compatibility (Comp vs Incomp) x Time interval (Short vs Long) x RTs–Bin (B1, B2, B3, B4) repeated measures ANOVAs and eventual significance of main effects of interaction was explored using Bonferroni post-hoc comparisons.

The visual inspection of scalp topographies revealed variations in the onset and peak times of the P300 and LPCt components as a function of time interval length and RTs-Bins. Consequently, individual amplitudes and latencies for these components were measured using a semi-automatic peak detection algorithm as implemented in Brain Analyzer 2.

In the case of P300, we considered a pool of central-posterior derivations (CP1, CP2, CPz, P1, Pz, P2; (Polich, 2007) during the 300-700 ms time window for short intervals and 180-360 ms time windows for long ones. Individual latency and amplitude data were entered in Space-Time Compatibility (Comp vs Incomp) x Time interval (Short vs Long) x RTs–Bin (B1, B2, B3, B4) repeated measures ANOVAs.

The LPCt was extracted from a pool of central-frontal sites (FC1, FCz, FC2, F1, Fz, F2; (Bannier et al., 2019; Bueno & Cravo, 2021; Gontier et al., 2009; Ofir & Landau, 2022; Paul et al., 2011) during a 600-1300 ms time window for Short intervals and 300-700 ms time window for Long ones. Individual amplitudes and latencies were entered in Space-Time Compatibility (Comp vs Incomp) x Time interval (Short vs Long) x RTs–Bin (B1, B2, B3, B4) repeated measures ANOVAs.

For all ANOVAs, significant main effects and interactions were further explored through Bonferroni post-hoc comparisons.

#### Analyses locked to target-offset: Lateralized Readiness Potential (C3 - C4)

As in the case of the analysis locked to target onset, for each of our experimental conditions, i.e., Space-Time Compatibility (Comp vs Incomp), Time Interval (Short vs Long) and RTs–Bin (B1, B2, B3, B4), we quantified the scalp motor regions’ differential activation by subtracting C4 from C3 EEG activity in a time window of -200 to 1500 ms. The statistical significance of these differential waveforms was calculated using a series of parametric continuous two-tailed one-sample T-tests (against zero) corrected for multiple comparisons (FDR) in the time domain, as implemented in Brainstorm.

*Analyses locked to manual responses: Bereitschaftspotential and Lateralized Readiness Potential (C3 > C4)*.

The Bereitschaftspotential (BP) was calculated as a) the mean amplitude from −600 ms to -200 ms at electrode Cz (Di Russo et al., 2017) for its negative subcomponent and b) the mean amplitude from −150 ms to -10 ms at the same Cz derivation for its pP1-2 complex subcomponent (Di Russo et al., 2017). Both components were analysed through a series of manual responses (Left, Right) x Space–Time Compatibility (Comp vs Incomp) x RTs–Bin (B1, B2, B3, B4) repeated measure ANOVAs. Significant main effects and interactions were explored through Bonferroni post-hoc comparisons.

The Lateralized Readiness Potential resulting from C3 > C4 differential activity was extracted from a -1200 + 300 ms time window and statistically assessed according to the same procedure described for onset- and offset-related epochs.

## Results

### Reaction times

We only found a significant STEARC at the slowest RTs in B3 and B4 (*Space-Time compatibility × RTs-bin interaction*: F(3, 84) = 5.3, p = .002, ƞp^2^ = .16; **Fig. 2**; B3: Comp: 493.7 ms vs Incomp: 510.1 ms, p = .02; B4: Comp: 673.4 ms vs Incomp 706.9 ms, p < .001;). We also found the typical Foreperiod Effect (ForEff) that is observed in STEARC tasks. In this case, once a time interval lasts more than a “short” one, participants are prepared to classify the interval as being “long” ahead of its offset (Niemi & Näätänen, 1981; Vallesi et al., 2011). This produces faster RTs to “long” than “short” intervals, i.e., the ForEff. In our study, the ForEff was highlighted by a significant *Time interval main effect* (F(1, 28) = 290.7, p < .001, ƞp^2^ = .9; **Fig. 2**) with faster RTs to long (427.1 ms) rather than short (557.6 ms) intervals. Generally speaking, the ForEff shows that in a STEARC task, participants can prepare in advance the response to the predictable end of “long” intervals, though not to that of “short” ones. With short intervals, the STEARC was significant at slower RTs in B3 and B4 (Space–Time Compatibility x RTs-bin interaction: F(3, 84) = 7.1, p = .0002, ƞp^2^ = .20; B3: Comp: 564.3 ms vs Incomp 590 ms, p < .001; B4: Comp: 727.6 ms vs Incomp 763.7 ms, p < .001). With long intervals, the STEARC was only found at the slowest RTs in B4 (Space–Time Compatibility x RTs-bin interaction (F(3, 84) = 2.6, p = .05, ƞp^2^ = .08; Comp: 619.1 ms vs Incomp 650.1 ms, p = .002; **Fig. 2**).

**Fig. 2:**
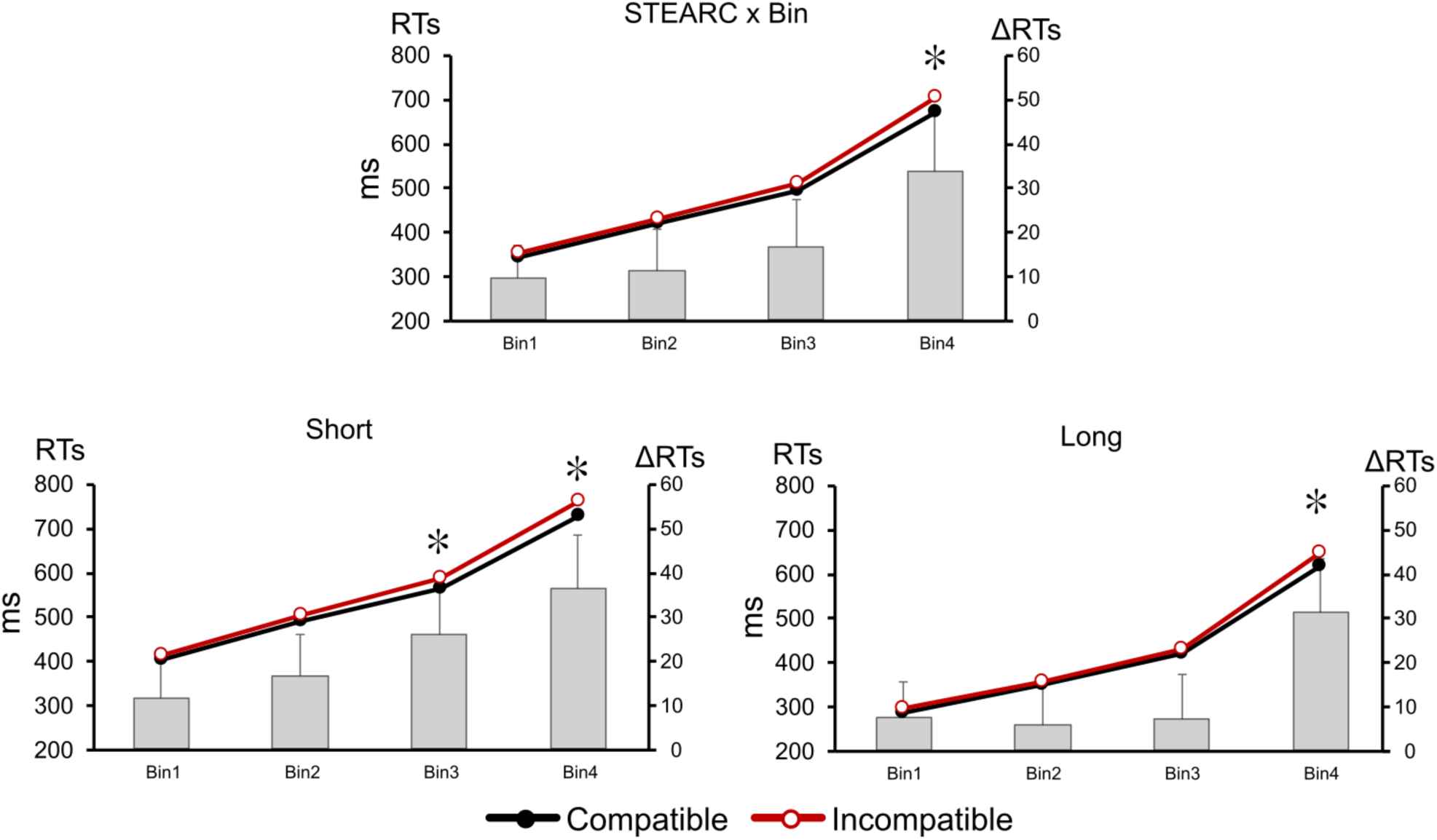
STEARC effect as a function of RTs speed (Short 1 sec. + Long 3 sec. intervals: Top panel); Bin 1 fastest RTs to Bin 4 slowest RTs. STEARC effects with Short (Bottom left panel) and Long (Bottom right panel) intervals. Left Y-axis: RTs in the Compatible (black line) and the Incompatible condition (red line). Right Y-axis: Incompatible minus Compatible RTs difference (grey bars). Significant differences are marked with an asterisk.

Response accuracy varied as a function of RTs speed (F (3, 84) = 18.935; p < 0.01; ƞp^2^ = 0.403). Bonferroni comparisons showed that errors increased significantly at the slowest RTs in B4 compared to all other bins (B1 = 0.83; B 2 = 0.43; B 3 = 0.7; B4 = 3; p < 0.01). A significant RTs-Bin × Time Interval interaction (F (3,84) = 65.569; p < 0.01; ƞp^2^ = 0.200) showed that this effect was driven by a decrease in response accuracy at B4 with long durations, while no change in accuracy was present with short durations across the different RT Bins (**Supplementary** Fig. 1 in Appendices). These results confirm that the STEARC, i.e., the spatial representation of elapsing time, takes place only when decisions on the length of time intervals are slow, though not when they are fast.

## EEG Results

### EEG activity locked to the onset of time intervals

#### The Contingent Negative Variation (CNV)

The CNV consists of a slow negative deflection in the EEG that occurs upon a first stimulus that foregoes a second task-relevant stimulus or action. In the case of time intervals, the onset of the time interval corresponds to the first stimulus and the offset to the second stimulus when the CNV resolves. The amplitude of the CNV that unfolds along a time interval was initially considered to reflect subjective timing (Kononowicz & Penney, 2016). Ensuing evidence has clarified that the CNV rather reflects the optimisation of cognitive and motor resources dedicated to interval timing, a process that is modulated by timekeeping mechanisms, though it does not exclusively reflect time processing (Kononowicz & Penney, 2016). During the late phase of the initial 1000 ms period of both short and long time intervals (550-990 ms), the CNV was significantly larger at faster RTs Bins (*RTs-Bin main effect*: F(3, 84) = 8.1, p < .001, ƞp^2^ = .22; B1: -2.16 µV vs B2: -1.99 µV, p = n.s.; **Fig. 3A** and **Fig. 3B**, see post-hoc comparisons in Supplementary Table 1). The same effect was also found in the late phase of long intervals (1700–2900 msec; *RTs-Bin main effect*: F(3, 84) = 4.5, p = .005, ƞp^2^ = .14; **Fig. 3B**, see post-hoc comparisons in Supplementary Table 2). These results show that in trials with slower RTs, when the STEARC occurs, during the presentation of time intervals the late component of the CNV is poorly developed.

**Fig. 3:**
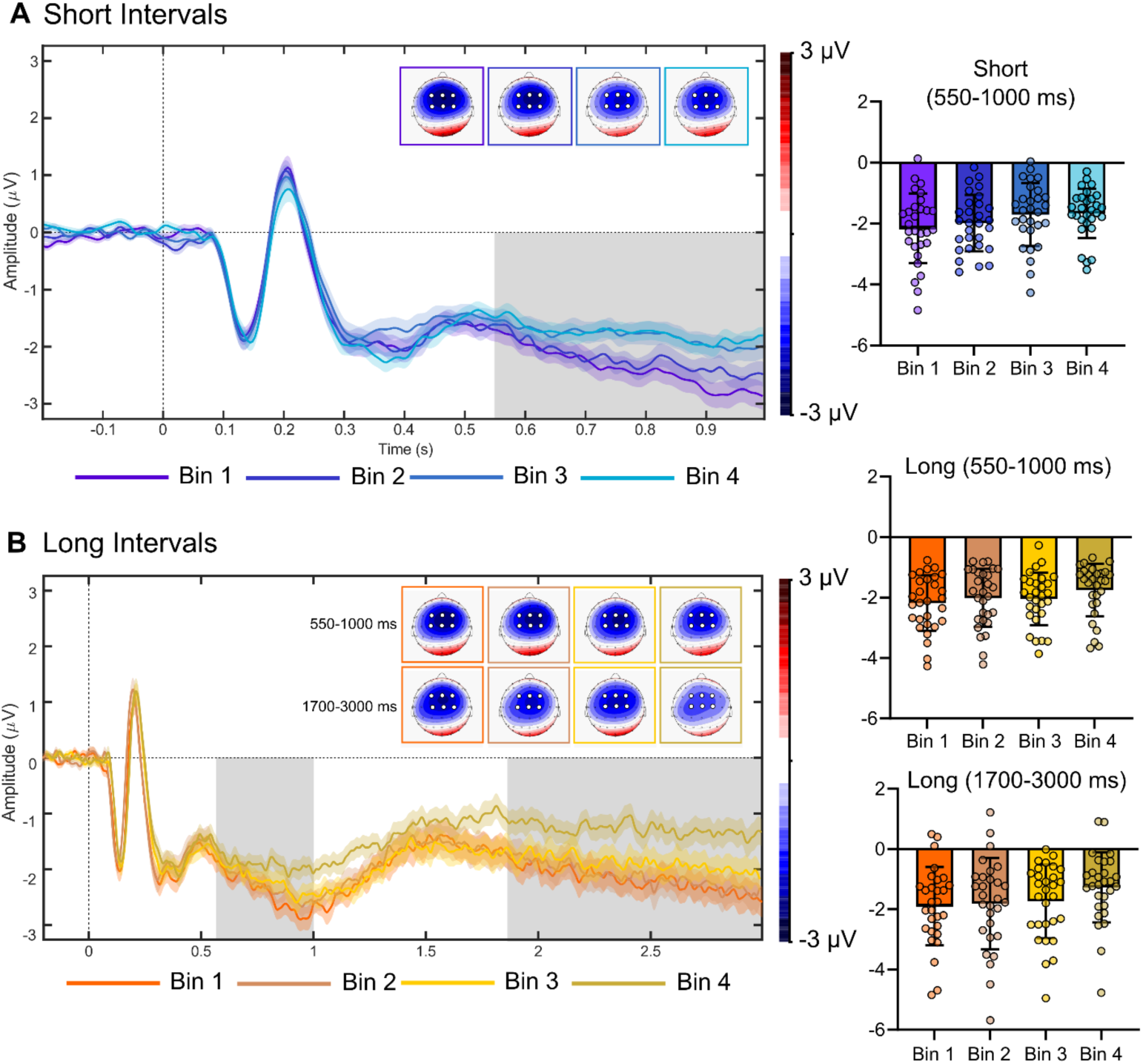
Waveforms, scalp topographies and individual/average amplitudes of the late CNV components locked to target-onset (time 0 on the x-axis). Time windows: 550-1000 ms, both Short and Long intervals, 1700-3000 ms only in Long intervals: The CNV is recorded over fronto-central derivations (FC1, FC2, FCz, C1, Cz, C2). Colour codes for RTs Bin 1, 2, 3, and 4 are reported in the figure. (A) Short 1 sec and (B) Long 3 sec intervals. Grey shade indicates periods of significant statistical differences between Bins.

#### C3-C4 differential activity (LRP): pre-activation of motor responses during the presentation of time intervals

We analysed the C3-C4 differential activity to explore whether during the presentation of the time interval the pre-activation of one hand response over the other changes as a function of the ensuing speed of response (see Vallesi et al., 2011): no significant differential C3 - C4 activity was found.

### EEG activity locked to the offset of time intervals

#### The N1/P2 complex: subjective timing

The amplitude of the N1/P2 component reflects subjective timing at the offset of time intervals when the CNV, which was developed during the same intervals, resolves (Kononowicz & Van Rijn, 2014). In addition, the amplitude of the N1/P2 component is also positively related to the difference between the subjective duration of a target interval and the duration of the reference to which it is compared (Kononowicz & Van Rijn, 2014). In our study the amplitude of N1 was modulated by the speed of RTs and the length of time intervals (F(3, 84) = 3.9, p = .01, ƞp^2^ = .11). With short intervals, the N1 was smaller at slower RTs in B4 (- 0.22 µV) as compared to faster RTs in B1 (-0.47 µV, p = .02) and B2 (-0.55 µV; p = .003). With long intervals, the N1 amplitude did not change as a function of RTs speed (**Fig. 4A and 4B**). The amplitude of the P2 was higher with short (1.27 µV) rather than long intervals (0.76 µV; F(1, 28) = 4.2, p = .04, ƞp^2^ = .13). A significant *Time interval x RTsBin interaction* (F(3, 84) = 3.1, p = .03, ƞp^2^ = .09 **Fig. 4A and 4B**) finally showed that with short intervals the amplitude of the P2 was larger at faster RTs in B1 (1.27 µV) as compared with all other Bins (B2: 1.21 µV, B3: 1.08 µV, B4: 1.16 µV; all p < .01). No similar difference was found with long intervals.

**Fig. 4:**
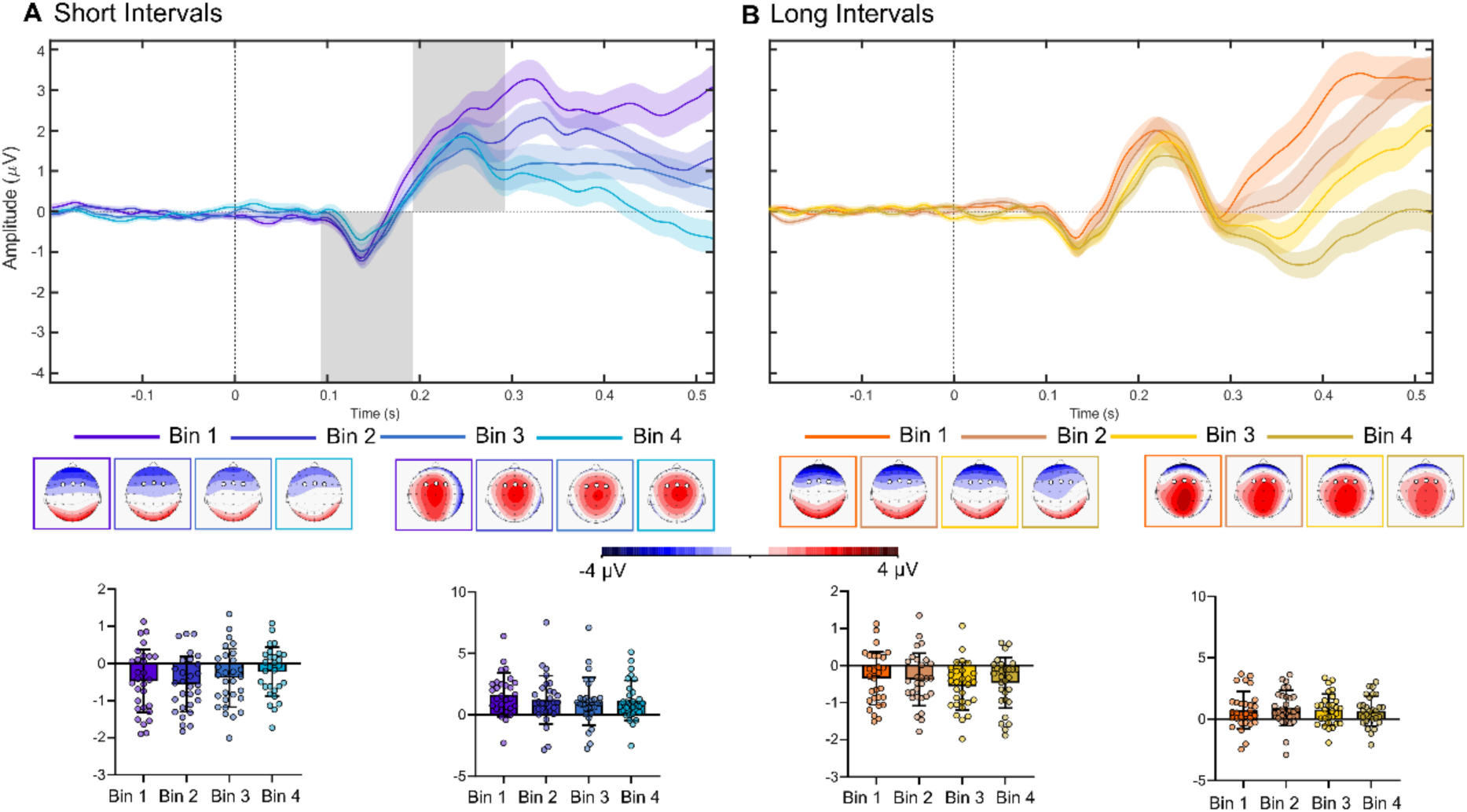
Waveforms, scalp topographies and individual/average amplitudes of the N1 (time window 90-180 ms) and P2 (time windows and 180-290 ms) components locked to target-offset (time 0 on the x-axis). Both components are recorded over frontal derivations (FC1, FC2, FCz). Colour codes for RTs Bin 1, 2, 3, and 4 are reported in the figure. (A) Short 1 sec and (B) Long 3 sec intervals. Grey shade indicates periods of significant statistical differences between Bins.

These results show that with short intervals, shallower N1 and P2 responses, i.e. less effective subjective timing, were associated with trials with slower RTs. The absence of a similar effect with long intervals suggests that, in this case, the ForEff helped predict the offset of these intervals and reduced the sensitivity of the N1 and P2 waves to subjective timing.

#### The P300: comparing time interval length

In timing tasks, the P300 marks the results of decision processes activated by the comparison between the target and the reference interval, so that the higher the difference between the two intervals, the higher the amplitude of the P300 (Ofir & Landau, 2022). In our study, the latency of the P300 was generally reduced with long as compared to short intervals (275.1 ms vs. 479.2 ms; *Time interval main effect*: F(1, 28) = 244.2, p < .001, ƞp^2^ = .89; **Fig. 5A**): this result was the likely consequence of the ForEff that allowed predicting the end of long intervals. With short intervals the latency increased at longer RTs (B1: 395 ms; B2: 458 ms; B3: 504 ms; B4: 561 ms; all p < .001; *Time interval x RTs Bin interaction*: F(3, 84) = 49.7, p < .001, ƞp^2^ = .63; **Fig. 5A**) while with long intervals no change in latency was found as a function of RTs speed (all p = n.s. **Fig. 5A**). The amplitude of the P300 was larger at the unpredictable offset of short intervals (5.76 µV) than at the predictable offset of long ones (3.79 µV; *Time interval main effect*: F(1, 28) = 26.7, p < .001, ƞp^2^ = .26; **Fig. 5A**), when it was larger at faster RTs in B1 (4.41 µV, p < .001) and B2 (3.96 µV, p < .001) than at slower RTs in B4 (3.19 µV; *Time interval x RTs Bin interaction*: F(3, 84) = 2.9, p = .04, ƞp^2^ = .09). These results show that in trials with slower RTs, when the STEARC emerged, there was and increase in P300 latency with short intervals and a reduction in P300 amplitude with long ones. These EEG markers suggest a slowed or less effective comparison between the duration of target intervals and that of the internal comparison reference.

**Fig. 5:**
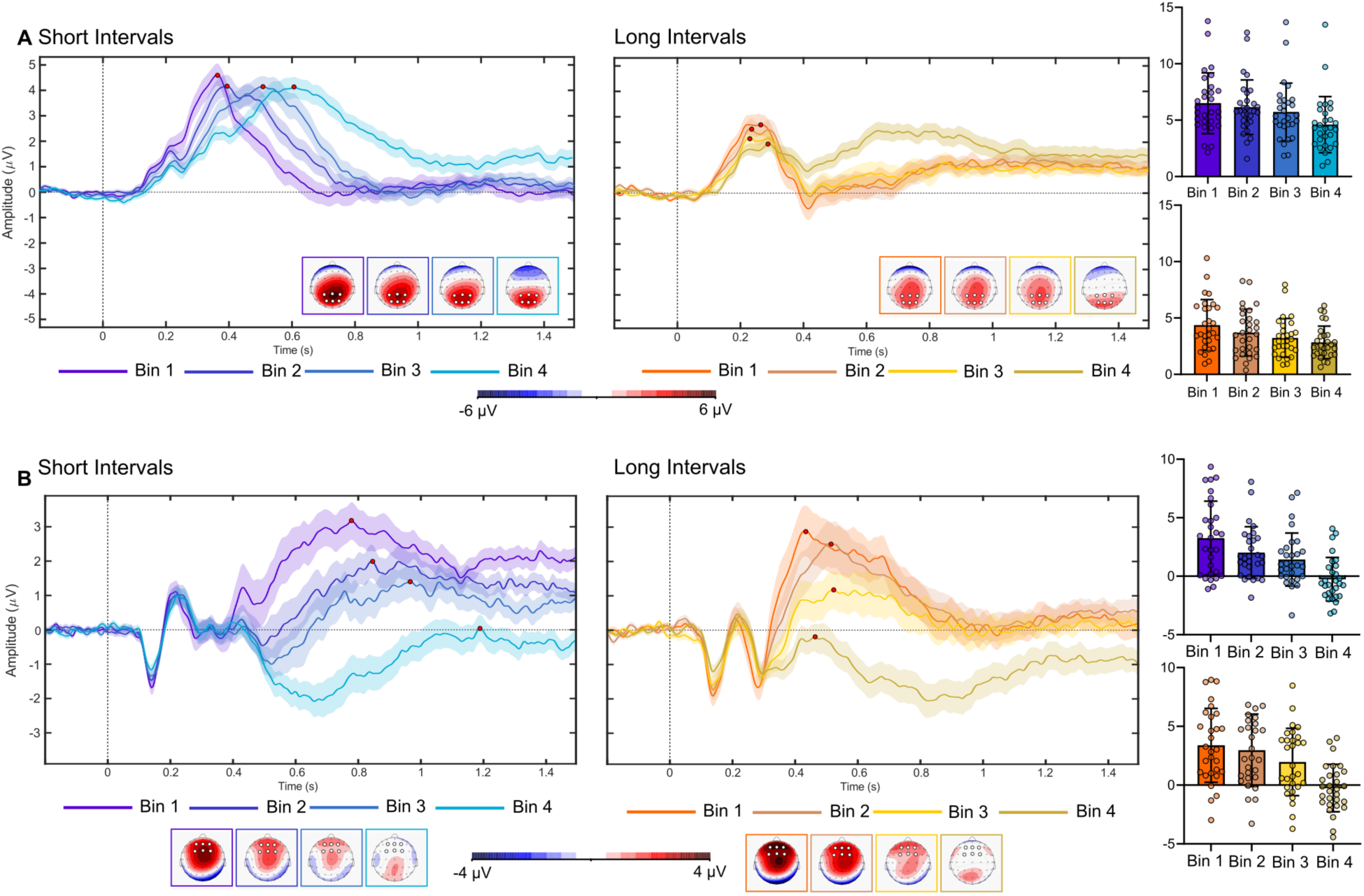
A) Waveforms, scalp topographies and individual/peak amplitudes of the P300 component locked to target-offset (time 0 on the x-axis), recorded over central-parietal derivations (CP1, CP2, CPz, P1, Pz, P2). Colour codes for RTs Bin 1, 2, 3, and 4 are reported in the figure. (A) Short 1 sec and (B) Long 3 sec intervals. Red dots indicate average peaks latency and amplitudes. **B)** Waveforms, scalp topographies and individual/peak amplitudes of the LPCt component locked to target-offset (time 0 on the x-axis), recorded over frontal derivations (FC1, FCz, FC2, F1, Fz, F2). Colour codes for RTs Bin 1, 2, 3, and 4 are reported in the figure. (A) Short 1 sec and (B) Long 3 sec intervals. Red dots indicate average peaks latency and amplitudes.

#### The Late Positive Component of timing (LPCt): accumulation of evidence favouring short vs long decisions

In timing tasks, the LPCt marks the accumulation of evidence toward “short” or “long” decision boundaries (Bannier et al., 2019; Bueno & Cravo, 2021; Kruijne et al., 2021; Ofir & Landau, 2022). As for the case of the P300, the ForEff reduced the latency of the LPCt with long (462.5 ms) as compared with short intervals (914.5 ms; *Time interval main effect*: F(1, 28) = 936.5, p < .001, ƞp^2^ = .92; **Fig. 5B**). In addition, while with long intervals the latency of the LPCt was equivalent across the different RTs Bins (all p = n.s. **Fig. 5B**), with short intervals it progressively increased as a function of RTs length (B1: 708 ms; B2: 851 ms; B3: 946 ms; B4: 1151 ms; all p < .03; *Time interval x RTs Bin interaction*: F(3, 84) = 41.1, p < .001, ƞp^2^ = .59; **Fig. 5B**). Finally, both with short and long intervals the amplitude of the LPCt was higher at faster RTs (B1: 3.32, B2: 2.49, B3: 1.69, B4: -0,25; all p < .001; *RTs Bin main effect*: F(3, 84) = 272.5, p < .001, ƞp^2^ = .7’; **Fig. 5B**). Similar to results with the P300 component, these findings show trials with slower RTs, when the STEARC occurs, were characterised by slowed or less effective accumulation of evidence in favour of short vs long decisions.

#### C3-C4 differential activity (LRP): pre-activation of motor responses at the offset of time intervals

The LRP locked to the offset of time intervals, captures the duration of the process that leads from the encoding of intervals to the selection of the duration response (Smulders & Miller, 2011). In our study, both with short Comp and Incomp intervals, an initial left-hand activation was observed 180-240 ms after target offset (**Fig. 6A**). In Incomp trials, e.g., “push the left button when the interval is short”, the initial left-hand activation was later replaced by right-hand activation 260-500 ms after target offset (**Fig. 6A**). With long Comp intervals, a right-hand activation was present 240-350 ms post-target offset (**Fig. 6A**). With long Incomp trials, significant left-hand activation was present 180-240 ms after target offset.

**Fig. 6:**
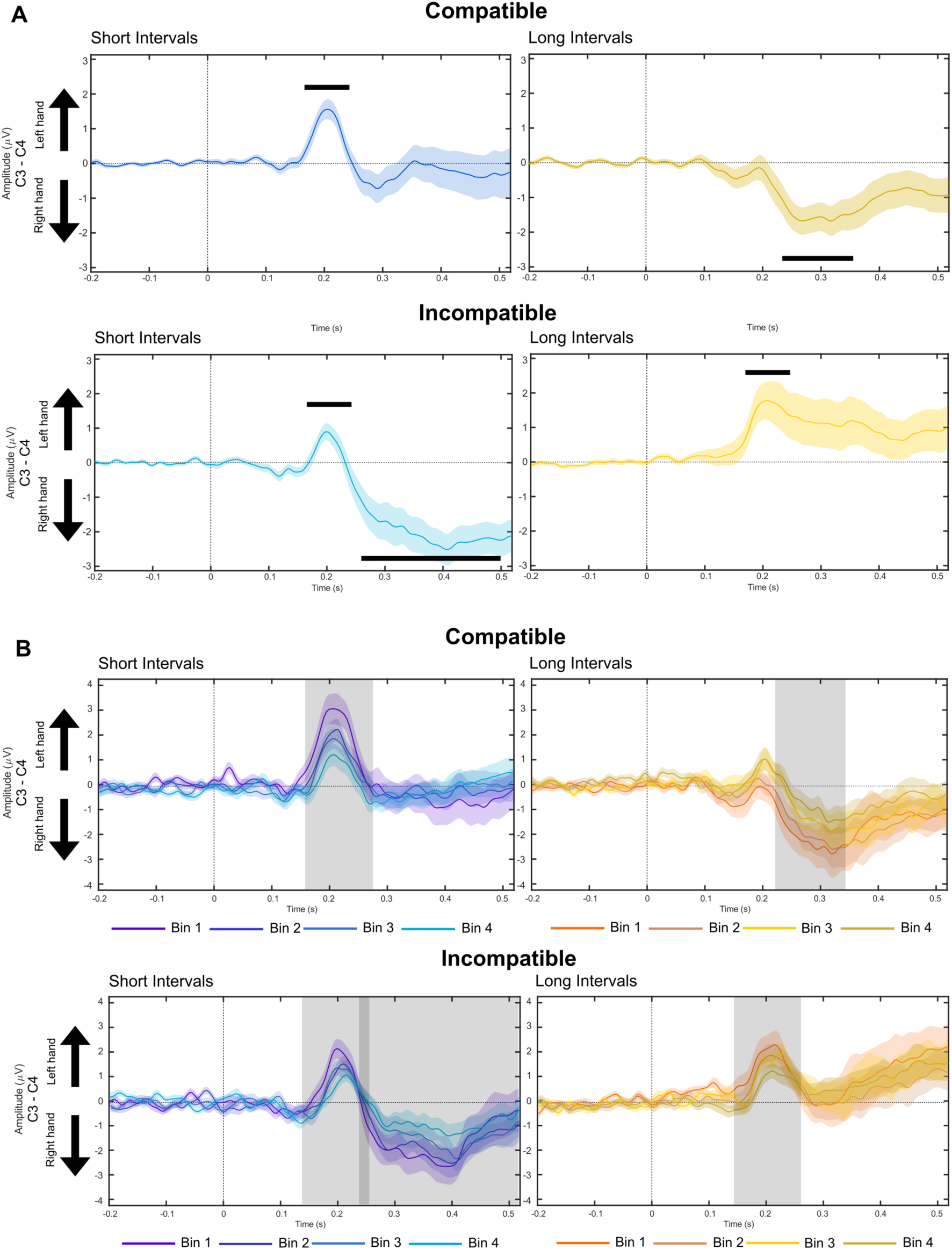
A) Differential C3 > C4 LRP waveforms locked to target-offset (time 0 on the x-axis) in the Compatible and Incompatible condition, with Short 1 sec (blue lines) and Long 3 sec (yellow lines) intervals. **B)** Differential C3 > C4 LRP waveforms locked to target-offset (time 0 on the x-axis) in the Compatible and Incompatible condition as a function of RTs-Bin for Short 1 sec (blue lines) and Long 3 sec (yellow lines) intervals. Colour codes for RTs Bin 1, 2, 3, and 4 are reported in the figure. Grey shades indicate periods of significant statistical differences between Bins (see also **Supplementary Tables 1 and 2**).

In addition, through a series of paired T-tests, we explored the amplitude of the LRP as a function of RT speed within time windows that showed significant differential C3-C4 activity (see Methods). In all types of trials, the pre-activation of the responding hand was reduced at slower RTs Bins (see **Fig. 6B** and **Supplementary Tables 3 and 4** in Appendices). This result shows that in the period ranging from the end of time intervals to the selection and activation of the motor response, trials with slower RTs, when the STEARC occurs, matched poorly defined motor pre-activation of the responding over the non-responding hand.

### EEG activity locked to manual responses

#### The Bereitschaftspotential (BP): response preparation and activation

The BP reflects cortical excitability linked to movement preparation and originates in the cingulate and supplementary motor cortex (Di Russo et al., 2017). The BP is characterised by an initial long leading negative component that starts 1-2 sec before the onset of a movement. This negative component is followed, 100-150 msec before movement onset, by a short pP2 positive wave that originates in the Anterior Insula (AIns) and reflects the accumulation of sensory evidence for categorising target stimuli (Boettiger & D’Esposito, 2005; for review, see Di Russo et al., 2017). In our experiment, the amplitude of the long leading negative component varied as a function of time interval length and RTs speed (F(3, 84) = 11.4, p < .001, ƞp^2^ = .36). With short intervals, it was enhanced at B1 (-1.2 µV) as compared to all other Bins (B2: -0.29 µV; B3: 0.16 µV; B4: 0.37 µV; all p < .0005). In seeming contrast, with long intervals, the same component was larger at slower RTs in B4 (-0.55 µV) compared to all other Bins (B1: -0.13 µV; B2: -0.04; µV B3: -0.02 µV; all p < .05; see **Fig. 7**). This seemingly paradoxical result was clarified by the analysis of the short pP2 sub-component of the BP. The pP2 was larger with short (1.4 µV) than long intervals (0.1 µV; F(1, 28) = 10.6, p = .002, ƞp^2^ = .27) and varied as a function of both interval length and RTs speed (F(3, 84) = 3.03, p = .03, ƞp^2^ = .09). With short intervals, it was selectively reduced at slower RTs in B4 (0.67 µV) compared to all other Bins (B1: 1.45 µV; B2: 1.69 µV; B3: 1.77 µV, all p < .03). With long intervals, the amplitude of the pP2 was smaller at B2 (0.41 µV) than at B1 (0.92 µV), became null at B3 (-0.11 µV) and at Bin 4 turned into a negative dip that by prolonging the foregoing long negative component caused its seemingly paradoxical enhancement in amplitude and duration (-1.16 µV; all p < .005; see **Fig. 7**). To summarise the analysis of the BO shows that trials with slower RTs when the STEARC occurs, match defective and late cortical excitability during motor response preparation.

**Fig. 7:**
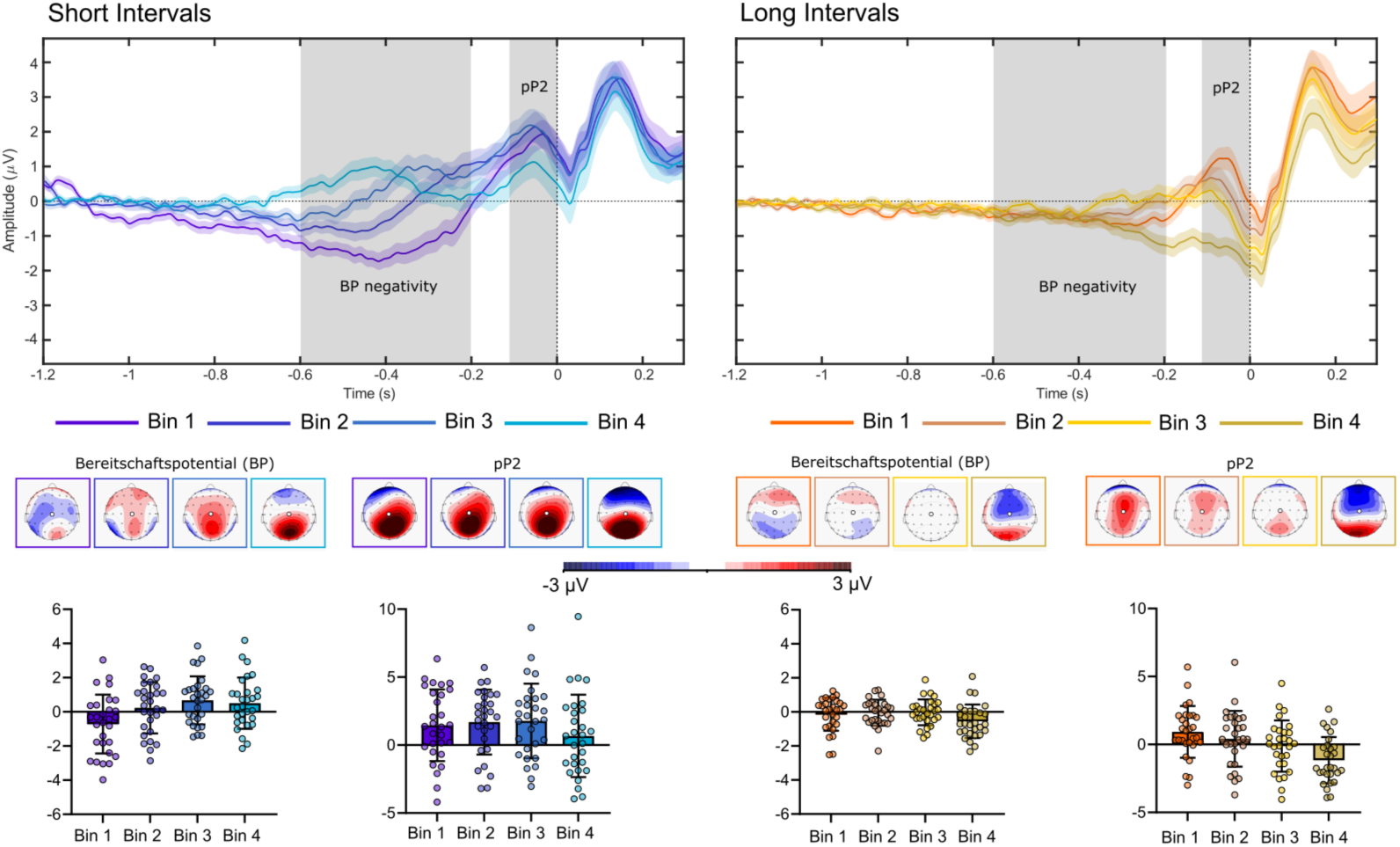
Waveforms, scalp topographies and individual/average amplitudes (time -600 -200 ms and -150 -10 ms; Cz derivation) of the negative and the pP1 component of the Bereitschaftspotential locked to manual response (time 0 on the x-axis) for Short (A) and Long (B) intervals. Colour codes for RTs Bin 1, 2, 3, and 4 are reported in the figure. Grey shade indicates periods of significant statistical differences between Bins.

#### C3-C4 differential activity (LRP): pre-activation of motor responses at response onset

The response-locked LRP wave reflects the duration of response activation and peripheral motor processes (Smulders & Miller, 2011). In short Comp trials, the activation of the left-hand started 170 ms before the response, while in long Incomp trials, it started 185 ms before the response (**Fig. 8A**). Compared to the performance of left-hand responses, right-hand responses showed a longer preparation. With long Comp trials, probably due to the foreperiod effect that allowed anticipating “long” duration decisions, the right-hand activation started around 280 ms before the response. In contrast, with Short incompatible trials, when the end of time intervals, in contrast with long trials, could not be anticipated, the same pre-activation was shorter, starting 210 ms before response (**Fig. 8A**).

**Fig. 8:**
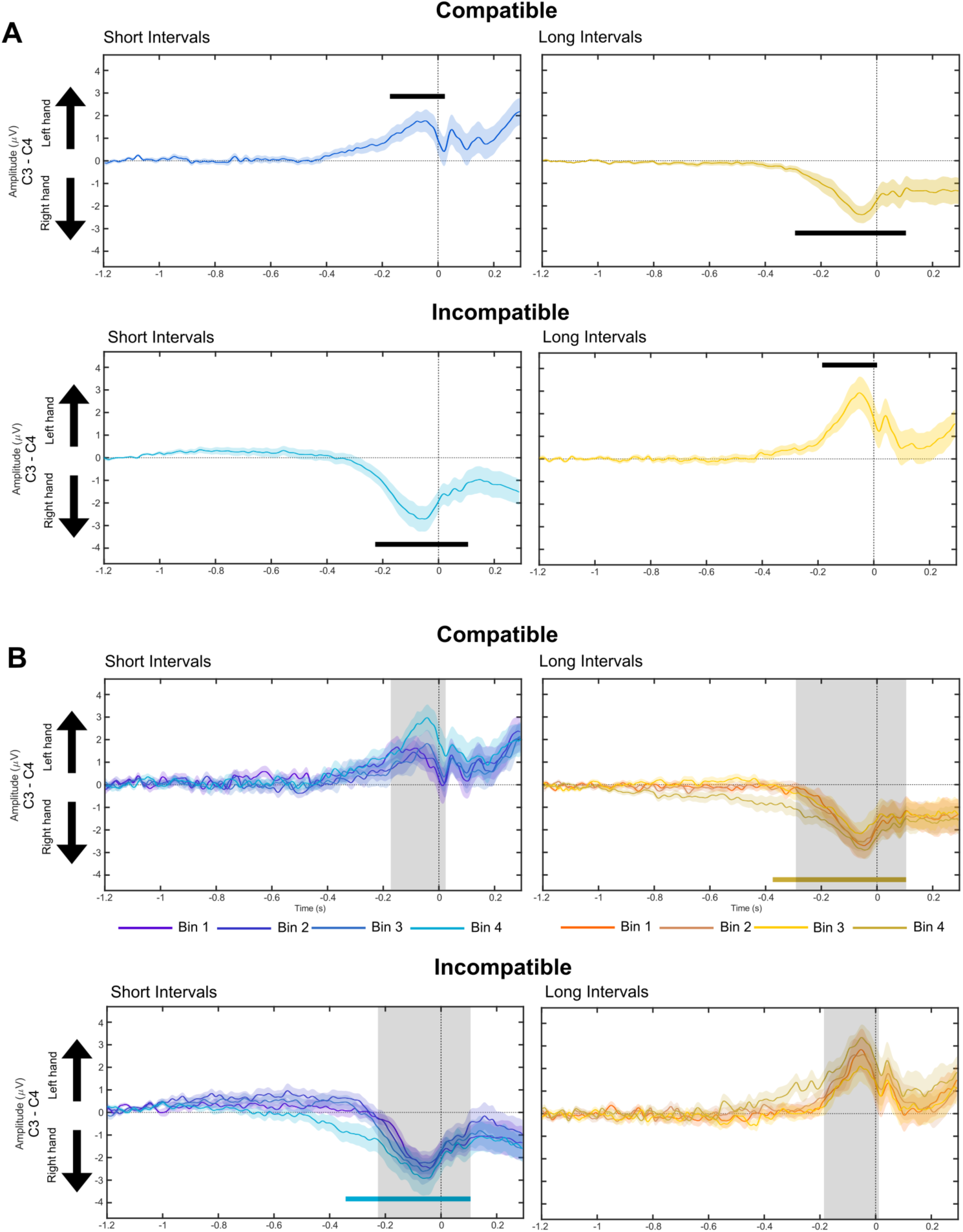
A) Differential C3 > C4 LRP waveforms locked to manual response (time 0 on the x-axis) in the Compatible and Incompatible condition with Short 1 sec (blue lines) and Long 3 sec (yellow lines) intervals. **B)** Differential C3 > C4 LRP waveforms locked to manual response (time 0 on the x-axis) in the Compatible and Incompatible condition, as a function of RTs Bin for Short 1 sec (blue lines) and Long 3 sec (yellow lines) intervals. Colour codes for RTs Bin 1, 2, 3, and 4 are reported in the figure. Grey shades indicate periods of significant statistical differences between Bins (see also Tables 1 and 2). Blue and yellow thick bars indicate periods of statistically significant C3 > C4 amplitude differences. Positive difference = left-hand more than right-hand pre-activation; Negative difference = right-hand more than left-hand pre-activation.

Using a series of t-tests, we then explored the LRP as a function of RTs speed within time windows showing significant C3-C4 differential activity (see Methods). With right-hand responses, the LRP started earlier at B4 than all other bins (**Fig. 8B**, top right and left bottom panels). With left-hand responses (**Fig. 8B**, top left and right bottom panels), the LRP was larger at B4 compared to all other Bins (see **Fig. 8B**). These results show that the occurrence of the STEARC at slower RTs was matched with larger and longer response-locked LRPs. This longer duration in response preparation compensates, in turn, for the poorly developed LRP that, in the same trials, was observed at the offset of time intervals (see previous result section).

## Discussion

Here, we show that the spatial representation of time revealed by the STEARC matches the modification of several EEG responses. The STEARC followed a cascade-like process that started from the encoding of time intervals. To begin with, shallower amplitudes of the late CNV phase that preceded the offset of time intervals were associated with the STEARC. With long time intervals, the CNV plateaued at the end of the first second, when the possible end of a short interval was expected, then faded and rose again up to the end of the interval. This biphasic trend suggests that the “short” 1-sec duration was taken as an anchor for the preparation of “long” decisions (Kononowicz & Penney, 2016; Ng et al., 2011). This confirms that rather than reflecting the accumulation of time pulses, in which case its amplitude should have increased linearly along the duration of long intervals, the CNV reflects optimisation of cognitive and resources ahead of interval offset (Kononowicz & Penney, 2016). Finally, no relationship was found as a function of RTs speed between the amplitude of the early N1P2 component of the CNV, which marks precision in initiating timing (Ng et al., 2011), and the amplitude of the later long negative component.

At the offset of short intervals, larger amplitudes of the N1P2 complex were associated with faster RTs and no STEARC. Since the N1/P2 reflects the perceived difference between the duration of a time interval and its comparison reference (Kononowicz & Van Rijn, 2014), this result shows that faster categorisation of short time intervals was associated with shortening of their subjective duration. In contrast, with long intervals, no amplitude difference in the N1P2 was found as a function of RTs speed. This latter finding was likely due to the ForEff that allowed participants to make “long” decisions ahead of interval offset and shows that strategic factors can reduce the amplitude and sensitivity of the N1P2 to subjective timing.

The late parietal P300 and a frontal Late Positive Component of Timing (LPCT) (for review, see Kononowicz et al., 2018) have been associated with evidence accumulation processes in determining the decision boundary for the length of time intervals. They are larger for intervals shorter than the boundary (Bannier et al., 2019; Giovannelli et al., 2014; Lindbergh & Kieffaber, 2013; Tarantino et al., 2010) because, in this case, the decision process must remain active up to the end of the interval (Ofir & Landau, 2022), and shallower for long intervals because, in this second case, decisions can be made ahead of the interval offset. The STEARC was associated with longer P300 latencies at short intervals and smaller amplitudes at long intervals. The P300 originates from a network of areas that includes as a relevant node the Temporal Parietal Junction (TPJ) (Igelström & Graziano, 2017; Polich, 2011), an area that signals the match/mismatch between expected and actual sensorimotor and cognitive events (Doricchi et al., 2010, 2022; Geng & Vossel, 2013). This suggests that in timing tasks, the P300 marks the subjective match/mismatch between a target interval and the comparison reference: with short intervals, when the STEARC is associated with longer P300 latencies, the release of match/mismatch signals is delayed, while with long intervals the ForEff smooths the relevance of match/mismatch signals at interval offset and, as a consequence, reduces the amplitude of the corresponding P300.

Both with short and long intervals, no LPCt was found at the slowest RTs when the STEARC emerged. This drop was associated with the late development of the P300 at short intervals and a sustained P300 activity at long ones (**Fig. 4**), just like if the development of the P300 interfered with the development of the ensuing LPCt. Based on the functional role assigned to the LCPt (Bannier et al., 2019; Bueno & Cravo, 2021; Kruijne et al., 2021; Ofir & Landau, 2022), its drop suggests a late and defective selection of S-R rules. This conclusion fits the finding that the STEARC was also associated with a late engagement of motor responses, as indicated by reduced or absent Bereitschaftspotentials (BP; Di Russo et al., 2017; Kornhuber & Deecke, 1985). This reduction affected both the long negative component of the BP, which originates in the cingulate and supplementary motor cortex and the later short pP2 component, which originates in the Anterior Insula (AntIns; for review, see Di Russo et al., 2017; Boettiger & D’Esposito, 2005).

The STEARC was previously associated with changes in the amplitude of the LRP, i.e. the C3-C4 differential activity, during and after the presentation of both short and long time intervals (Vallesi et al., 2011). On this ground, a likely expectation would be that the larger the LRP, the stronger the STEARC. In complete contrast with this prediction, we have found that at the end of time intervals and in correspondence with the progressive building up of the STEARC at slower RTs, the amplitude of the LRP reflecting the activation of Compatible responses decreased, rather than increased, with respect to the amplitude of the LRP reflecting Incompatible responses (**Fig. 9**). This result points out that when the preparation of the Compatible responding hand is effective and the response of the non-responding hand well inhibited, responses are faster and the STEARC does not take place.

**Fig. 9:**
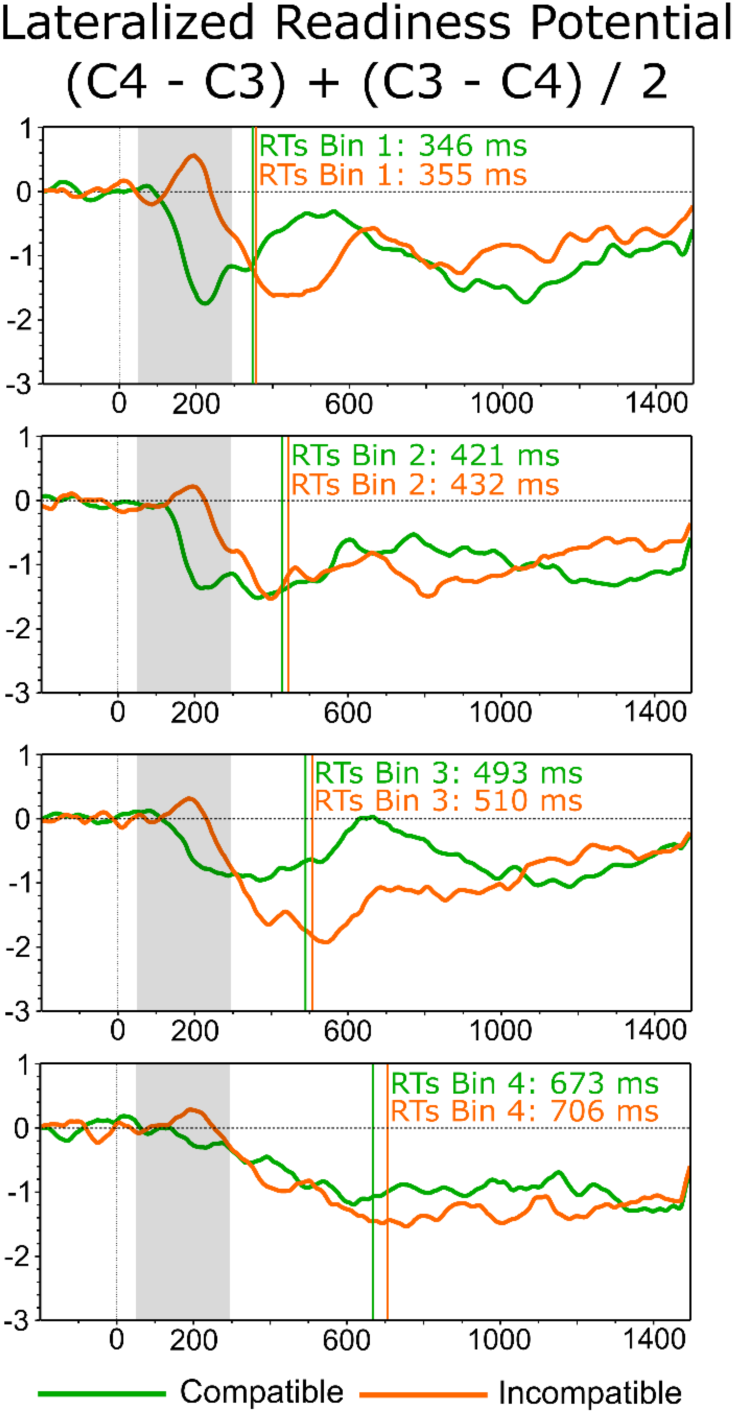
Differential C3 > C4 LRP waveforms averaged between the left and right hand and locked to the offset of time intervals (time 0 on the x-axis) in the Compatible (green lines) and Incompatible (orange lines) condition at Bin 1 (top row) to 4 (bottom row). Grey shades indicate periods of significant statistical differences between. Vertical bars indicate the average manual RTs at each Bin.

### A unifying account of the EEG correlates of the STEARC

How do we interpret the multifaceted EEG correlates of the STEARC? A unifying causal mechanism can be identified in the role that the catecholamine system, i.e. dopamine (DA) and norepinephrine (NE), plays in motivation, cognitive control and timekeeping (Fung et al., 2021; Silvetti et al., 2018). Modelling studies (Silvetti et al., 2018, 2023) suggest that a loop between the medial prefrontal cortex (MPFC) and brainstem nuclei optimizes catecholamine release as a function of task demands, motivation and expected value of cognitive engagement. Optimization is based on learning to control internal variables influenced by catecholamines, such as effort, neural plasticity and attention, i.e. meta-learning (Silvetti et al., 2018). According to these models, during the STEARC task, enhanced control of signals from MPFC to the catecholamine brainstem nuclei results in higher DA and NE release. This improves the encoding of time intervals and speeds up decisions on their length as well as response selection, resulting in shorter RTs. Importantly, optimisation of catecholamine release is a stochastic process, which means that, holding all environmental variables constant, optimization fluctuates from trial-to-trial determining variability in RTs and EEG responses. These fluctuations are reflected by the CNV that originates in the MPFC, the neural module that drives optimization of meta-learning (Silvetti et al., 2018: **Fig. 10**). To summarize, this model sketches a causal link among the trial-by-trial variability of the MPFC control signal, the variability of catecholamine release and that of RTs.

**Fig. 10:**
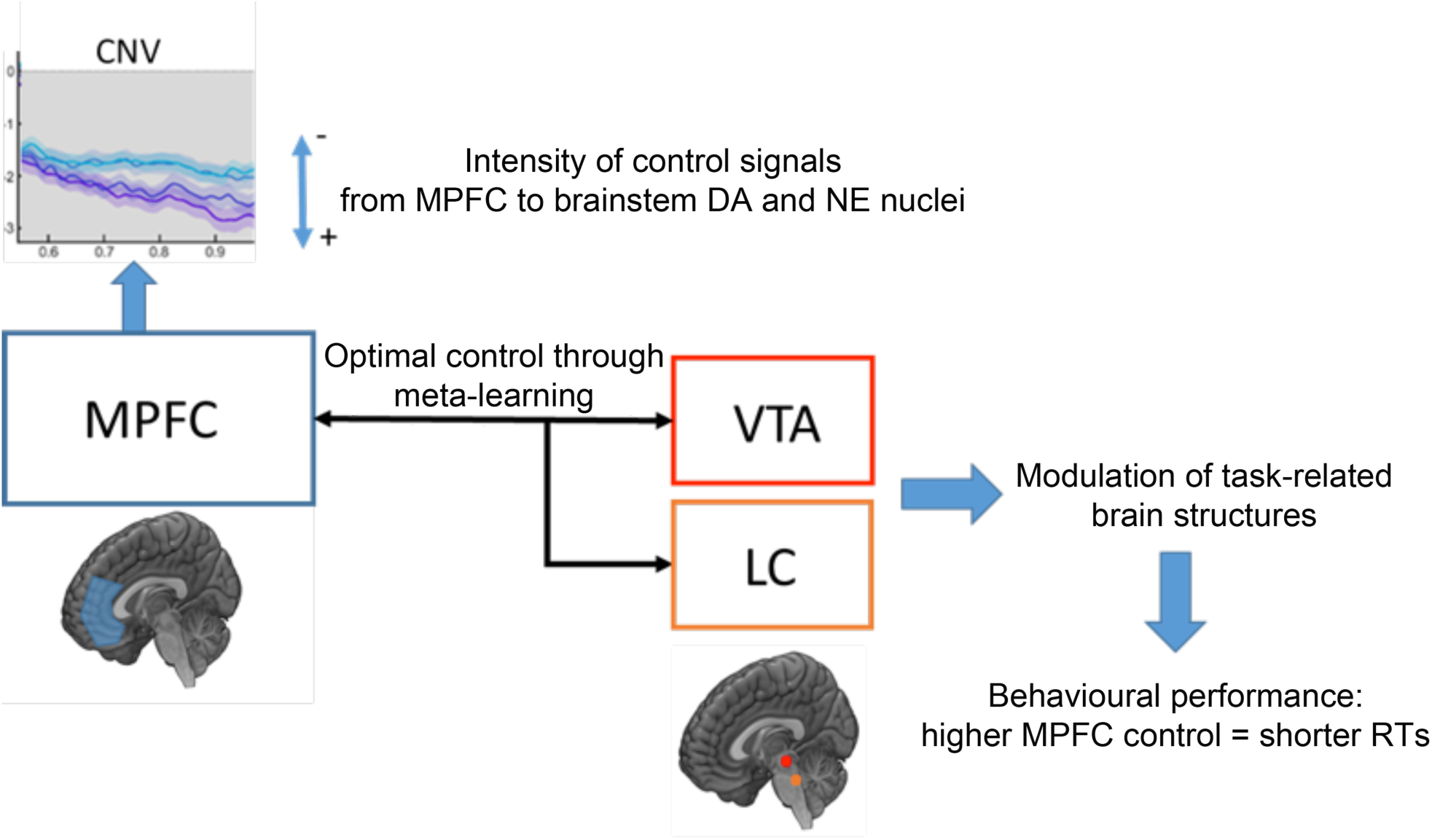
Model of the MPFC-catecholamine system and its role in behavioural performance. MPFC (Medial Prefrontal Cortex), VTA (Ventral Tegmental Area, DA transmitter), LC (Locus Coeruleus, NE transmitter), DA (Dopamine), NE (Norepinephrine).

In line with this interpretation, we found that the initial marker of the STEARC was a drop in CNV amplitude, i.e. the EEG signature of the drop of control signal from MPFC to the brainstem (**Fig. 10**). Since levels of DA correlate with CNV amplitude (Kononowicz & Penney, 2016; Linssen et al., 2011; Tecce et al., 1975), it can be concluded that phasic drops in DA activity and poor optimisation of cognitive resources characterised trials with shallower CNV, slow RTs and a significant STEARC. Drops in DA also slow down the accumulation of time pulses, leading to underestimation of time durations (Fung et al., 2021). Our study found that slower RTs and the STEARC were associated with reduced N1/P2 complex amplitude, which indexes the subjective duration of time intervals (Kononowicz & Van Rijn, 2014). This suggests that the drop in DA activity during the encoding CNV-phase, affected the perceived duration of time intervals reflected by the N1/P2 component. Finally, DA and NE also interact in the modulation of P300 components, i.e. P3A and P3b (Polich, 2007; Warren et al., 2023). In both cases, lower DA or NE activity reduces P300 amplitude, while effects on corresponding latencies are currently unclear (see [59] for review). In our study, the STEARC was associated with longer latency (short intervals) and reduced amplitude (long intervals) of the P300: also, these effects are compatible with the hypothesis that the STEARC matches reduced catecholaminergic activity.

## Conclusions

Cultural practices in reading, writing, and inspecting the environment impose ordinal and spatial sequencing of sensorimotor and cognitive events both during the encoding of a series of temporal events and the learning of number series (Buzsáki & Llinás, 2017; Pitt & Casasanto, 2020). These learning episodes enhance the brain’s representation of time and numbers by incorporating visual-spatial features. The history of human cultures and science offers many examples of how visual-spatial representations of time and numbers have helped and shaped humans’ capacity to comprehend, interpret, and make sense of the environment (Buonomano & Rovelli, 2021; Buzsáki & Llinás, 2017; Giaquinto, 2011). The “curving of space-time” in general relativity is a famous example of these spatially informed conceptualisations (Buonomano & Rovelli, 2021). Nonetheless, recent evidence shows that the mental arrangement of consecutive time ticks along a left-to-right organised Mental Time Line and numbers along a Mental Number Line are not inherent to the brain representation of time or numbers. These arrangements are rather triggered by the use of spatial response codes in the task at hand (Pinto et al., 2019; Pinto, Pellegrino, Lasaponara et al., 2021; Pinto, Pellegrino, Marson et al., 2021; Scozia et al., 2023), by the availability of lateralised spatial affordances, e.g., left and right arm waving to indicate the past and the future, or because of the presence of operational symbols, e.g. “+” or “-“, that re-activate overlearned left-to-right mental representations of addition and subtraction operations (see (Bonato et al., 2021) for operations run on time durations). The results of the present study reinforce this conclusion by showing that even when left/right spatial response codes are used to classify the length of time intervals, a corresponding left-to-right mental representation of time is bypassed when timing decisions are fast. This finding provides a new insight into the brain mechanisms that subtend the widespread use of the mental spatial representation of the flow of time and, possibly, a new tool for studying separately the non-spatial and spatial brain mechanisms of timekeeping.

## Acknowledgements.

This work was supported by a grant PRIN 2022 (Code: 2022WPY4L3) from the Italian Ministry of Research and Education (MUR) to F.D. We thank Juan David Banor Salazar from the Universidad de San Buenaventura Medellin (Colombia) for help in data collection.

## Acknowledgements

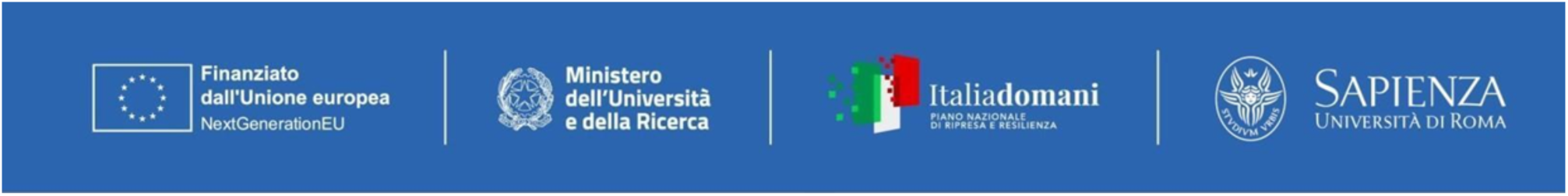

## Appendices

**Supplementarty Figure 1.**
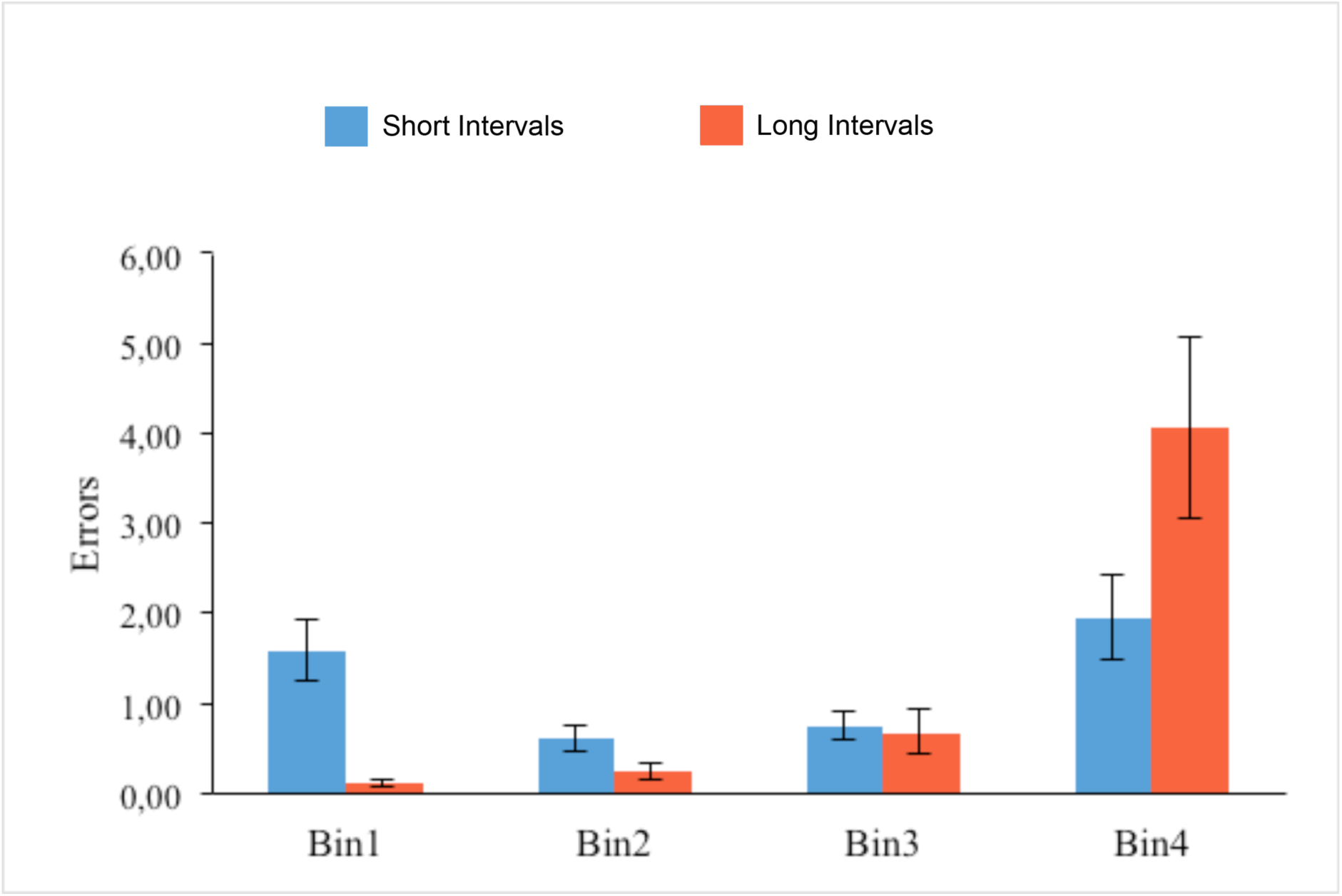
Performance accuracy in the STEARC task, i.e. number of errors, as a function of time interval length.

**Supplementary Table 1:**
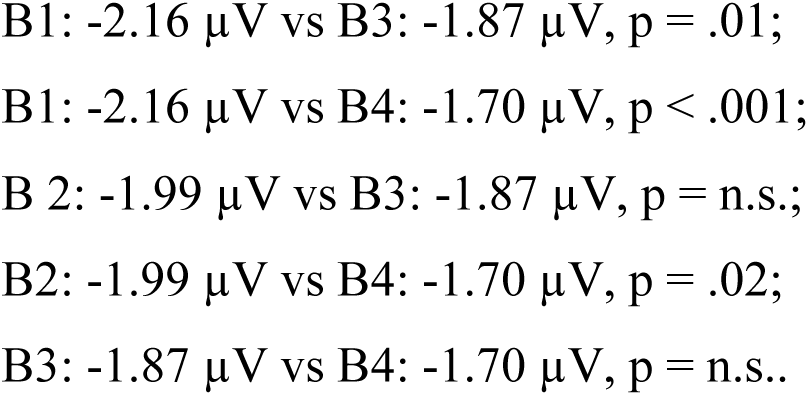
CNV amplitude during the initial 1000 ms of time intervals Bin by Bin post hoc comparisons:

**Supplementary Table 2:**
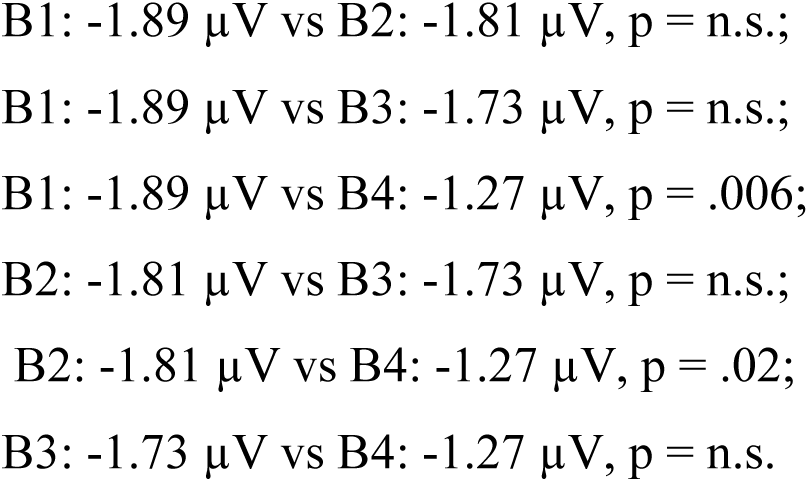
CNV amplitude during the late phase of long time intervals Bin by Bin post hoc comparisons:

**Supplementary Table 3:**
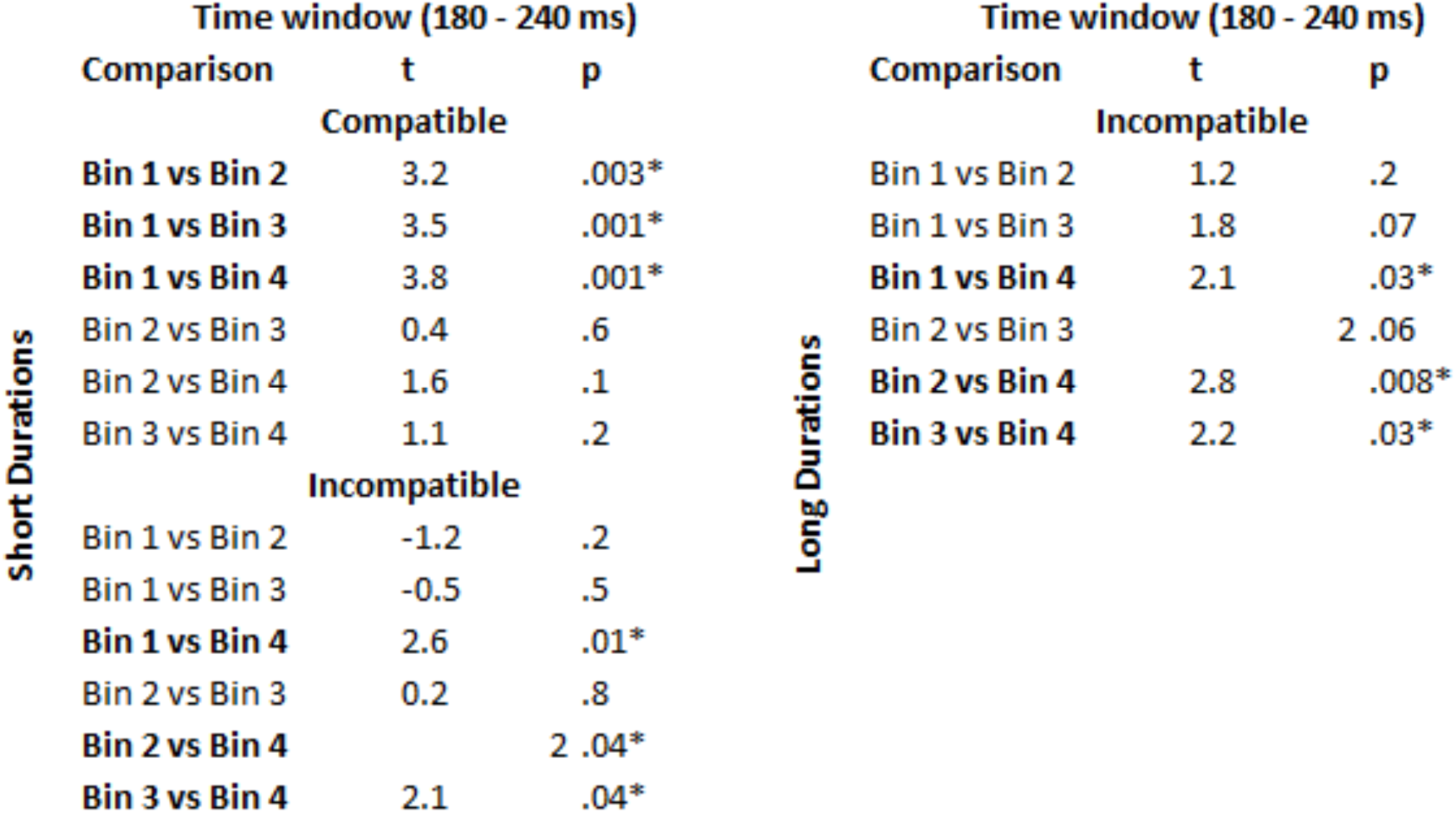
T-tests comparisons of the C3 > C4 positive differential activity found at 180-240 ms in Compatible and Incompatible Short and Long-interval conditions.

**Supplementary Table 4:**
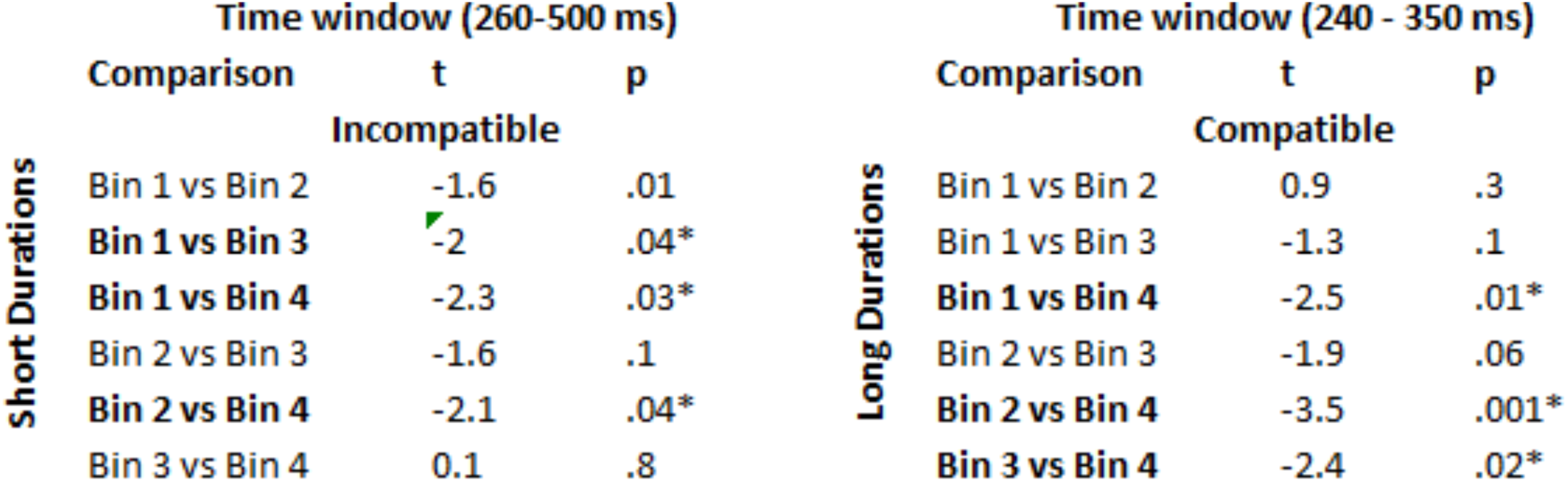
T-tests comparisons of the C3 > C4 negative differential activity found at 260-350 ms and 260-500 ms in Compatible and Incompatible conditions of Short and Long intervals.

## References.

1. Bannier, D., Wearden, J., Le Dantec, C. C., & Rebaï, M. (2019). Differences in the temporal processing between identification and categorization of durations: A behavioral and ERP study. Behavioural Brain Research, 356, 197–203. 10.1016/j.bbr.2018.08.027

2. Boettiger, C. A., & D’Esposito, M. (2005). Frontal Networks for Learning and Executing Arbitrary Stimulus-Response Associations. The Journal of Neuroscience, 25(10), 2723– 2732. 10.1523/JNEUROSCI.3697-04.2005

3. Bonato, M., D’Ovidio, U., Fias, W., & Zorzi, M. (2021). A momentum effect in temporal arithmetic. Cognition, 206, 104488. 10.1016/j.cognition.2020.104488

4. Bonato, M., Zorzi, M., & Umiltà, C. (2012). When time is space: Evidence for a mental time line. Neuroscience & Biobehavioral Reviews, 36(10), 2257–2273. 10.1016/j.neubiorev.2012.08.007

5. Boroditsky, L., Fuhrman, O., & McCormick, K. (2011). Do English and Mandarin speakers think about time differently? Cognition, 118(1), 123–129. 10.1016/j.cognition.2010.09.010

6. Brunia, C. H. M., & van Boxtel, G. J. M. (2000). Motor preparation. In J. T. Caccioppo, L. G. Tassinary, & G. G. Berntson (A c. Di), Handbook of Psychophysiology. 2nd edition (pp. 507–532). Cambridge University Press.

7. Bueno, F. D., & Cravo, A. M. (2021). Post-interval EEG activity is related to task-goals in temporal discrimination. PLOS ONE, 16(9), e0257378. 10.1371/journal.pone.0257378

8. Buonomano, D., & Rovelli, C. (2021). *Bridging the neuroscience and physics of time* (Versione 1). arXiv. 10.48550/ARXIV.2110.01976

9. Buzsáki, G., & Llinás, R. (2017). Space and time in the brain. Science, 358(6362), 482–485. 10.1126/science.aan8869

10. Callizo-Romero, C., Tutnjević, S., Pandza, M., Ouellet, M., Kranjec, A., Ilić, S., Gu, Y., Göksun, T., Chahboun, S., Casasanto, D., & Santiago, J. (2020). Temporal focus and time spatialization across cultures. Psychonomic Bulletin & Review, 27(6), 1247–1258. 10.3758/s13423-020-01760-5

11. Casasanto, D., & Boroditsky, L. (2008). Time in the mind: Using space to think about time. Cognition, 106(2), 579–593. 10.1016/j.cognition.2007.03.004

12. Conson, M., Cinque, F., Barbarulo, A. M., & Trojano, L. (2008). A common processing system for duration, order and spatial information: Evidence from a time estimation task. Experimental Brain Research, 187(2), 267–274. 10.1007/s00221-008-1300-5

13. Di Russo, F., Berchicci, M., Bozzacchi, C., Perri, R. L., Pitzalis, S., & Spinelli, D. (2017). Beyond the “Bereitschaftspotential”: Action preparation behind cognitive functions. Neuroscience & Biobehavioral Reviews, 78, 57–81. 10.1016/j.neubiorev.2017.04.019

14. Doricchi, F., Lasaponara, S., Pazzaglia, M., & Silvetti, M. (2022). Left and right temporal-parietal junctions (TPJs) as “match/mismatch” hedonic machines: A unifying account of TPJ function. Physics of Life Reviews, 42, 56–92. 10.1016/j.plrev.2022.07.001

15. Doricchi, F., Macci, E., Silvetti, M., & Macaluso, E. (2010). Neural correlates of the spatial and expectancy components of endogenous and stimulus-driven orienting of attention in the Posner task. Cerebral Cortex, 20(7), 1574–1585.

16. Faul, F., Erdfelder, E., Lang, A.-G., & Buchner, A. (2007). G*Power 3: A flexible statistical power analysis program for the social, behavioral, and biomedical sciences. Behavior Research Methods, 39(2), 175–191. 10.3758/BF03193146

17. Fuhrman, O., McCormick, K., Chen, E., Jiang, H., Shu, D., Mao, S., & Boroditsky, L. (2011). How Linguistic and Cultural Forces Shape Conceptions of Time: English and Mandarin Time in 3D. Cognitive Science, 35(7), 1305–1328. 10.1111/j.1551-6709.2011.01193.x

18. Fung, B. J., Sutlief, E., & Hussain Shuler, M. G. (2021). Dopamine and the interdependency of time perception and reward. Neuroscience & Biobehavioral Reviews, 125, 380–391. 10.1016/j.neubiorev.2021.02.030

19. Geng, J. J., & Vossel, S. (2013). Re-evaluating the role of TPJ in attentional control: Contextual updating? Neuroscience & Biobehavioral Reviews, 37(10), 2608–2620. 10.1016/j.neubiorev.2013.08.010

20. Giaquinto, M. (2011). Visual thinking in mathematics: An epistemological study (1. publ. in paperback). Oxford Univ. Press.

21. Giovannelli, F., Ragazzoni, A., Battista, D., Tarantino, V., Del Sordo, E., Marzi, T., Zaccara, G., Avanzini, G., Viggiano, M. P., & Cincotta, M. (2014). “…the times they aren’t a-changin’…” rTMS does not affect basic mechanisms of temporal discrimination: A pilot study with ERPs. Neuroscience, 278, 302–312. 10.1016/j.neuroscience.2014.08.024

22. Gontier, E., Paul, I., Le Dantec, C., Pouthas, V., Jean-Marie, G., Bernard, C., Lalonde, R., & Rebaï, M. (2009). ERPs in anterior and posterior regions associated with duration and size discriminations. Neuropsychology, 23(5), 668–678. 10.1037/a0015757

23. Igelström, K. M., & Graziano, M. S. A. (2017). The inferior parietal lobule and temporoparietal junction: A network perspective. Neuropsychologia, 105, 70–83. 10.1016/j.neuropsychologia.2017.01.001

24. Ishihara, M., Keller, P., Rossetti, Y., & Prinz, W. (2008). Horizontal spatial representations of time: Evidence for the STEARC effect. Cortex, 44(4), 454–461. 10.1016/j.cortex.2007.08.010

25. Kononowicz, T. W., & Penney, T. B. (2016). The contingent negative variation (CNV): Timing isn’t everything. Current Opinion in Behavioral Sciences, 8, 231–237. 10.1016/j.cobeha.2016.02.022

26. Kononowicz, T. W., & Van Rijn, H. (2014). Decoupling Interval Timing and Climbing Neural Activity: A Dissociation between CNV and N1P2 Amplitudes. The Journal of Neuroscience, 34(8), 2931–2939. 10.1523/JNEUROSCI.2523-13.2014

27. Kononowicz, T. W., Van Rijn, H., & Meck, W. H. (2018). Timing and Time Perception: A Critical Review of Neural Timing Signatures Before, During, and After the To-Be-Timed Interval. In J. T. Wixted (A c. Di), *Stevens’* Handbook of Experimental Psychology and Cognitive Neuroscience (1a ed., pp. 1–38). Wiley. 10.1002/9781119170174.epcn114

28. Kornhuber, H. H., & Deecke, L. (1985). The starting function of the SMA. Behavioral and Brain Sciences, 8(4), 591–592. 10.1017/S0140525X00045210

29. Kruijne, W., Olivers, C. N. L., & Van Rijn, H. (2021). Neural Repetition Suppression Modulates Time Perception: Evidence From Electrophysiology and Pupillometry. Journal of Cognitive Neuroscience, 33(7), 1230–1252. 10.1162/jocn_a_01705

30. Lindbergh, C. A., & Kieffaber, P. D. (2013). The neural correlates of temporal judgments in the duration bisection task. Neuropsychologia, 51(2), 191–196. 10.1016/j.neuropsychologia.2012.09.001

31. Linssen, A. M. W., Vuurman, E. F. P. M., Sambeth, A., Nave, S., Spooren, W., Vargas, G., Santarelli, L., & Riedel, W. J. (2011). Contingent negative variation as a dopaminergic biomarker: Evidence from dose-related effects of methylphenidate. Psychopharmacology, 218(3), 533–542. 10.1007/s00213-011-2345-x

32. Ng, K. K., Tobin, S., & Penney, T. B. (2011). Temporal Accumulation and Decision Processes in the Duration Bisection Task Revealed by Contingent Negative Variation. Frontiers in Integrative Neuroscience, 5. 10.3389/fnint.2011.00077

33. Niemi, P., & Näätänen, R. (1981). Foreperiod and simple reaction time. Psychological Bulletin, 89(1), 133–162. 10.1037/0033-2909.89.1.133

34. Núñez, R. E., & Sweetser, E. (2006). With the Future Behind Them: Convergent Evidence From Aymara Language and Gesture in the Crosslinguistic Comparison of Spatial Construals of Time. Cognitive Science, 30(3), 401–450. 10.1207/s15516709cog0000_62

35. Ofir, N., & Landau, A. N. (2022). Neural signatures of evidence accumulation in temporal decisions. Current Biology, 32(18), 4093–4100.e6. 10.1016/j.cub.2022.08.006

36. Ouellet, M., Santiago, J., Israeli, Z., & Gabay, S. (2010). Is the Future the Right Time? Experimental Psychology, 57(4), 308–314. 10.1027/1618-3169/a000036

37. Paul, I., Wearden, J., Bannier, D., Gontier, E., Le Dantec, C., & Rebaï, M. (2011). Making decisions about time: Event-related potentials and judgements about the equality of durations. Biological Psychology, 88(1), 94–103. 10.1016/j.biopsycho.2011.06.013

38. Pinto, M., Pellegrino, M., Lasaponara, S., Scozia, G., D’Onofrio, M., Raffa, G., Nigro, S., Arnaud, C. R., Tomaiuolo, F., & Doricchi, F. (2021). Number space is made by response space: Evidence from left spatial neglect. Neuropsychologia, 154, 107773.

39. Pinto, M., Pellegrino, M., Marson, F., Lasaponara, S., Cestari, V., D’Onofrio, M., & Doricchi, F. (2021). How to trigger and keep stable directional Space–Number Associations (SNAs). Cortex, 134, 253–264.

40. Pinto, M., Pellegrino, M., Marson, F., Lasaponara, S., & Doricchi, F. (2019). Reconstructing the origins of the space-number association: Spatial and number-magnitude codes must be used jointly to elicit spatially organised mental number lines. Cognition, 190, 143–156.

41. Pitt, B., & Casasanto, D. (2020). The correlations in experience principle: How culture shapes concepts of time and number. Journal of Experimental Psychology: General, 149(6), 1048–1070. 10.1037/xge0000696

42. Polich, J. (2007). Updating P300: An integrative theory of P3a and P3b. Clinical Neurophysiology, 118(10), 2128–2148.

43. Polich, J. (2011). Neuropsychology of P300. Oxford University Press. 10.1093/oxfordhb/9780195374148.013.0089

44. Ratcliff, R. (1979). Group reaction time distributions and an analysis of distribution statistics. Psychological Bulletin, 86(3), 446–461. 10.1037/0033-2909.86.3.446

45. Rubichi, S., Nicoletti, R., Iani, C., & Umiltà, C. (1997). The Simon effect occurs relative to the direction of an attention shift. Journal of Experimental Psychology: Human Perception and Performance, 23(5), 1353–1364. 10.1037/0096-1523.23.5.1353

46. Scozia, G., Pinto, M., Lozito, S., Lasaponara, S., Binetti, N., Pazzaglia, M., & Doricchi, F. (2023).Space is a late heuristic of elapsing time: New evidence from the STEARC effect. Cortex, 164, 21–32. 10.1016/j.cortex.2023.03.009

47. Shibasaki, H., Barrett, G., Halliday, E., & Halliday, A. M. (1981). Cortical potentials associated with voluntary foot movement in man. Electroencephalography and Clinical Neurophysiology, 52(6), 507–516. 10.1016/0013-4694(81)91426-7

48. Silvetti, M., Lasaponara, S., Daddaoua, N., Horan, M., & Gottlieb, J. (2023). A Reinforcement Meta-Learning framework of executive function and information demand. Neural Networks, 157, 103–113. 10.1016/j.neunet.2022.10.004

49. Silvetti, M., Vassena, E., Abrahamse, E., & Verguts, T. (2018). Dorsal anterior cingulate-brainstem ensemble as a reinforcement meta-learner. PLOS Computational Biology, 14(8), e1006370. 10.1371/journal.pcbi.1006370

50. Smulders, F. T. Y., & Miller, J. O. (2011). The Lateralized Readiness Potential. Oxford University Press. 10.1093/oxfordhb/9780195374148.013.0115

51. Tadel, F., Baillet, S., Mosher, J. C., Pantazis, D., & Leahy, R. M. (2011). Brainstorm: A User-Friendly Application for MEG/EEG Analysis. Computational Intelligence and Neuroscience, 2011, 1–13. 10.1155/2011/879716

52. Tarantino, V., Ehlis, A.-C., Baehne, C., Boreatti-Huemmer, A., Jacob, C., Bisiacchi, P., & Fallgatter, A. J. (2010). The time course of temporal discrimination: An ERP study. Clinical Neurophysiology, 121(1), 43–52. 10.1016/j.clinph.2009.09.014

53. Tecce, J. J., Cole, J. O., & Savignano-Bowman, J. (1975). Chlorpromazine effects on brain activity (contingent negative variation) and reaction time in normal women. Psychopharmacologia, 43(3), 293–295. 10.1007/BF00429268

54. Teghil, A., Marc, I. B., & Boccia, M. (2021). Mental representation of autobiographical memories along the sagittal mental timeline: Evidence from spatiotemporal interference. Psychonomic Bulletin & Review, 28(4), 1327–1335. 10.3758/s13423-021-01906-z

55. Toma, K., Mima, T., Matsuoka, T., Gerloff, C., Ohnishi, T., Koshy, B., Andres, F., & Hallett, M. (2002). Movement Rate Effect on Activation and Functional Coupling of Motor Cortical Areas. Journal of Neurophysiology, 88(6), 3377–3385. 10.1152/jn.00281.2002

56. Vallesi, A., Binns, M. A., & Shallice, T. (2008). An effect of spatial–temporal association of response codes: Understanding the cognitive representations of time. Cognition, 107(2), 501–527. 10.1016/j.cognition.2007.10.011

57. Vallesi, A., McIntosh, A. R., & Stuss, D. T. (2011). How time modulates spatial responses. Cortex, 47(2), 148–156. 10.1016/j.cortex.2009.09.005

58. Vallesi, A., Weisblatt, Y., Semenza, C., & Shaki, S. (2014). Cultural modulations of space–time compatibility effects. Psychonomic Bulletin & Review, 21(3), 666–669. 10.3758/s13423-013-0540-y

59. Warren, C. V., Kroll, C. F., & Kopp, B. (2023). Dopaminergic and norepinephrinergic modulation of endogenous event-related potentials: A systematic review and meta-analysis. Neuroscience & Biobehavioral Reviews, 151, 105221. 10.1016/j.neubiorev.2023.105221

